# Enzyme fluctuations data improve inference of metabolic interaction networks with an inverse differential Jacobian approach

**DOI:** 10.1101/2023.12.11.570118

**Authors:** Jiahang Li, Wolfram Weckwerth, Steffen Waldherr

## Abstract

The development of next-generation sequencing and single-cell technology has generated vast genome-scale multi-omics datasets. Dedicated mathematical algorithms are required to dissect intricate molecular causality within metabolic networks using these datasets. Based on the network analysis, recent research has introduced the inverse differential Jacobian algorithm, which combines metabolic interaction network construction and covariance matrix analysis of genome-scale metabolomics data to elucidate system regulatory factors near steady-state dynamics. Traditionally, these studies assumed metabolomics variations solely resulted from metabolic system fluctuations, acting independently on each metabolite. However, emerging evidence highlights the role of internal network fluctuations, particularly from the gene expression fluctuations, leading to correlated perturbations on metabolites.

In this article, we propose a novel approach that exploits these correlations to reconstruct relevant metabolic interactions. Thereby, enzymes exhibiting significant variances in activity values serve as indicators of large fluctuations in their catalyzed reactions. By integrating this information in an inverse Jacobian algorithm, we are able to exploit the underlying reaction network structure to improve the construction of the fluctuation matrix required in the inverse Jacobian algorithm. After a comprehensive assessment of three critical factors affecting the algorithm’s accuracy, we conclude that using the enzyme fluctuation data significantly enhances the inverse Jacobian algorithm’s performance. We applied this approach to a breast cancer dataset with two different cell lines, which highlighted metabolic interactions where fluctuations in enzyme gene expression yield a relevant difference between the cell lines.

## Introduction

With the advancement of next-generation sequencing and single-cell technology [1], a plethora of genome-scale OMICS datasets has become available. One primary focus of systems biology is the exploration of interactions among the components of biological systems and the mechanisms through which these interactions influence the function and behavior of the system [2, 3]. For instance, the investigation of how enzymes and metabolites interact within a metabolic pathway remains an ongoing question in the field of metabolomics [4-9]. From a systematic perspective, metabolic networks typically involve many non-linear interactions between metabolites and enzymes where perturbations on each compound play a crucial role [10-12]. Current systematic approaches for metabolomics data analysis can be divided into three categories: statistical analysis, dynamic modeling, and network analysis. Statistical methods, particularly machine learning algorithms provide valuable insights into cellular activities under various treatment conditions and identify key biomarkers within the biological system [13, 14]. These methods encompass techniques such as Principal Components Analysis (PCA) [15], clustering analysis [16], deep learning [17], genetic algorithm [18] and boosting machine learning methods [19, 20]. However, they lack the ability to detect perturbation sites associated with dynamics of the underlying metabolic network system. On this aspect, kinetic models can be constructed to improve our systemic insight into a metabolic network. Over the last two decades, extensive biological studies have developed manually curated or optimized kinetic models which are available in databases such as the BioModels Database [21].

To analyze dynamic regulations of the system, it is natural to build kinetic models fitting the metabolomics measurements. During this modeling process, kinetic parameters can be obtained from previous models [22] or estimated from experimental data. Overall, the modelling process is a long and iterative endeavor involving literature mining and parameter tuning, and sometimes it does not yield satisfactory results [23, 24]. Moreover, constructing a comprehensive biological model often necessitates time-series data, which is not always available due to experimental constraints. Thus, the kinetic modeling approach is primarily limited to small-scale models. To our knowledge, there is only one genome-scale kinetic modeling work in literature [25].

Another mathematical approach uses network analysis, often beginning with the examination of a correlation or covariance matrix estimated from the measurements [9, 10, 26]. Subsequent studies aim to uncover underlying causal relationships from the correlation network [12, 27]. The available comprehensive information about metabolic network structure is now being collected in different databases [28-30], and can be used as extra information for the interaction network inference [31, 32]. This approach to network inference relies on stochastic fluctuations, which induce variability in metabolite concentrations that can be used to infer metabolic interactions even in steady state with an inverse Jacobian algorithm [26]. In recent years, several studies have in addition developed inverse differential Jacobian algorithms, which provide a convenient way to infer differences in the dynamics of metabolic networks between different conditions from metabolomics data [6, 31-37].

In these studies, the fluctuations are claimed to independently act on all metabolites [26, 31, 32, 35, 38]. Independent fluctuations would primarily arise from stochastically perturbed transport between the studied system and its external environments. However, more recent studies have provided evidence that fluctuations also originate from within the network itself [39-45]. Over the last years, researches have shown that fluctuations in the gene regulatory network result in variations in enzyme activities, either do to variations in enzyme expression or in post-translational regulation, further leading to variations in the reaction rate parameters [42, 44, 45]. Due to the reaction network structure, this leads to correlated fluctuations acting on multiple metabolites. More specifically, we define a fluctuation matrix D to be used in the inverse Jacobian algorithm, which in former algorithms was assumed to be diagonal, while the fluctuations we consider here give rise to a non-diagonal structure. Moreover, in contrast to previous approaches, we reconstruct the fluctuation matrix D from the variance of the enzyme activities. Figure 1 presents the work scheme of this approach. By utilizing the reaction network information in the structure of both the Jacobian J and the fluctuation matrix D, we can improve the regression-loss based inverse differential Jacobian algorithm.

**Figure 1.**
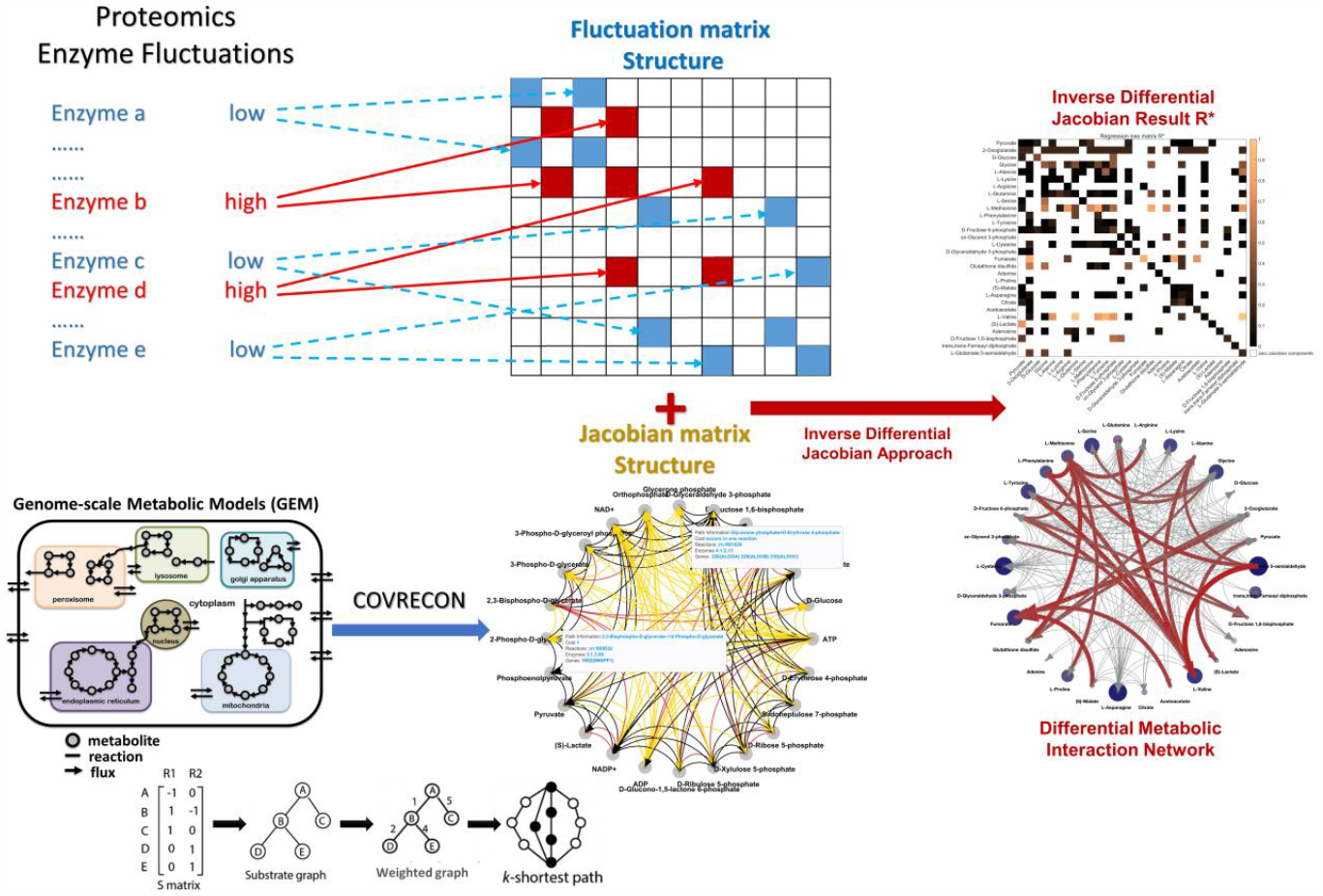
Work scheme of the inverse Jacobian analysis with a non-diagonal fluctuation matrix.

We evaluate this approach with several published models collected from the EBI BioModels database [21], and also apply the approach to a breast cancer dataset with two different conditions. We compared the inverse Jacobian results with different assumptions on the structure of the fluctuation matrix D [46]. We show that the inverse differential Jacobian approach is enhanced by incorporating this information. However, off-diagonal elements pertaining to small fluctuations can be disregarded without significant drawback. Furthermore, we analyzed the precision and accuracy of our algorithm. This analysis is based on varying three factors: (1) the amplitude of fluctuations, (2) the number of off-diagonal components in the fluctuation matrix D, and (3) the fluctuation magnitude of off-diagonal components compared to the diagonal components in the fluctuation matrix D. The main findings are that the inverse differential Jacobian algorithm is significantly enhanced using the non-diagonal structure with an integrative sampling; nevertheless, it remains feasible to assume a diagonal structure of D in the inverse differential Jacobian algorithm when the number of off-diagonal fluctuations is large or their magnitude is relatively small. In conclusion, this article gives the first comprehensive analysis of the impact of non-diagonal fluctuations on the inverse Jacobian approach. Furthermore, we introduced an enhanced inverse Jacobian algorithm provided with structure information on the fluctuation matrix. Furthermore, we show how to infer the structure of the fluctuation matrix from covariance data of enzyme activities. This approach holds significant promise for improving our understanding of metabolic network dynamics and the robustness of the inverse Jacobian algorithm in various applications.

## Methods

### The differential Jacobian

Consider a biological system that consists of n compounds (metabolites, proteins) denoted by{X_i_}_i=1… n_. The vector M = {M_i_} = {|X_i_|} represent the concentrations of the n compounds. The system dynamics can be modeled with the set of ordinary differential equations (ODEs):

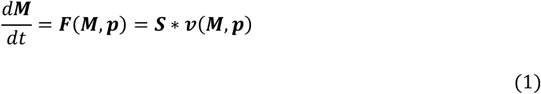

where ***S*** is the stoichiometry matrix formed by the stoichiometric coefficients of all the m reactions ***v***(***M, p***) = [*v*_1_(***M, p***), …, *v*_*m*_(***M, p***)] in the system. The reaction rates *v*_*i*_(***M, p***), *i* = 1, …, *m* are usually modelled by Michaelis-Menten kinetics [47] or mass action equations [48] depending on metabolite concentrations ***M*** and parameters (e.g., enzyme activities) ***p***.

The steady-state Jacobian matrix ***J*** of Equation (1) is defined as a *R*^*n*×*n*^ matrix in which *J*_*ij*_ is the first-order derivative of the rate of change *f*_*i*_ of the metabolite concentration *M*_*j*_, evaluated at steady state, noted as 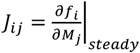 :

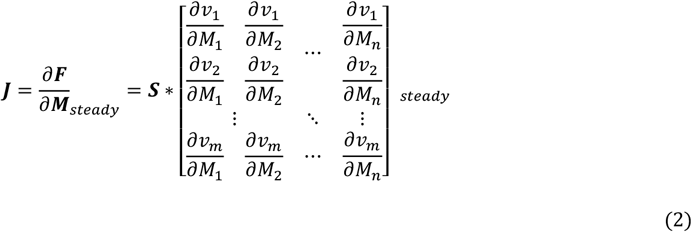

In a previous study, Steuer et al. [26] used a model perturbed by stochastic fluctuations to establish a relation between the covariance matrix *C* of the metabolite concentrations and the steady-state Jacobian matrix of the system ***J*** given by the so-called Lyapunov equation

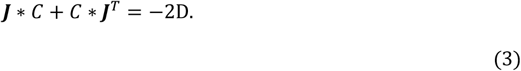

Thereby, *C* ∈ *R*^*n*×*n*^ is the covariance matrix of the metabolite concentrations *M*_*j*_ around their steady-state values 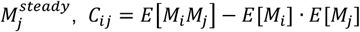, and the fluctuation matrix D is the covariance of fluctuations sources acting on the system dynamics.

A typical inverse task in this setup is to infer the Jacobian matrix J, representing the interactions in the network, from estimates for C and potentially D from steady-state metabolomics data. Furthermore, for the inverse differential Jacobian, we more specifically infer the difference between the Jacobian matrices for two biological conditions, for example a healthy and disease condition, abbreviated as ‘h’ and ‘d’, thereby identifying regulations in biochemical interactions that act differently between these two conditions. The differential Jacobian *D****J*** is defined as [31]:

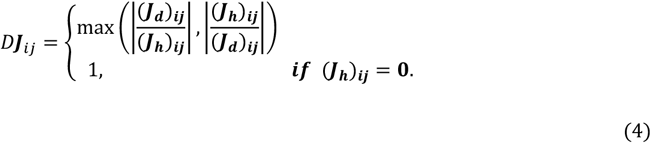

with the two conditional Jacobian matrix at steady-state denoted as ***J***_***h***_ and ***J***_***d***_.

The initial problem in the inverse task is that the system is under-determined, due to that covariance matrix C and fluctuation matrix D are symmetric matrixes but the Jacobian matrix ***J*** is in general not symmetric. Sun and Weckwerth addressed this problem by constraining the structure of the inferred Jacobian matrix from a topological metabolic interaction network, yielding entries which are constrained to zero in the Jacobian matrix ***J*** [32]. They argued that the topological network can be built from genome-scale network reconstruction and based on public accessible database, such as KEGG [29] and BioCyc [49]. At that time, the proposed algorithm was limited by two issues: they rely on structural network information that needs to be assembled manually, and they encounter numerical instability problem due to ill-conditioned regression problems for large-scale metabolic networks [31, 36]. In a more recent study, we developed a fully automated COVRECON workflow and related Matlab toolbox [31] that can automatically construct the topology of the metabolic interaction network from KEGG [29], BiGG [28] and ModelSeed [30] databases, and replaces the ill-conditioned regression problem with a regression loss-based inverse Jacobian algorithm.

### Inferring the fluctuation matrix structure from enzyme activity data

Previous inverse Jacobian algorithms [26, 31, 32, 35, 36] have assumed that independent stochastic noise affects each metabolite individually, giving rise to a diagonal fluctuation matrix D in the Lyapunov equation (3). In this article, we introduce additional stochastic fluctuations in enzyme activity [39, 40, 42, 43]. Through the coupling of metabolites by reactions, these result in correlated fluctuations of metabolite concentrations. Importantly, we assume that noise acting on the enzyme activity is statistically independent of noise acting directly on metabolites.

Suppose in the same system with dynamics Equation (1), the steady state concentration of *X*_*i*_ is 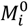, when the parameters ***p*** take the nominal values ***p***_**0**_. We consider stochastic perturbations on the metabolites as Gaussian noise vector ***ω***, and stochastic perturbations of parameters ***p*** with a Gaussian noise vector ***τ***. Thus, the reaction parameter vector becomes ***p***(*t*) = ***p***_**0**_ + ***τ***. Small fluctuations of metabolite concentrations around steady state are described by

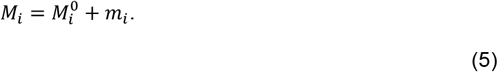

Overall, the dynamics with stochastic fluctuations thus become

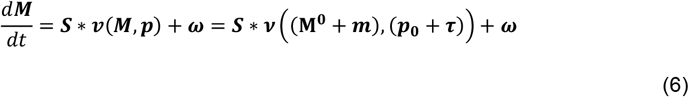

A Taylor expansion near the system’s nominal steady state (**M**^**0**^, ***p***_**0**_) then yields

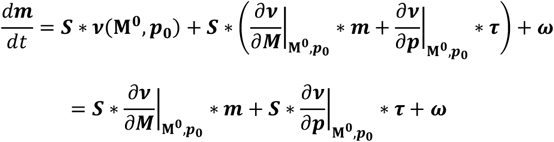

Observing that 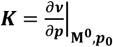 is a constant matrix, we get

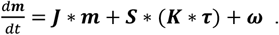

Thus the fluctuation matrix D is determined as,

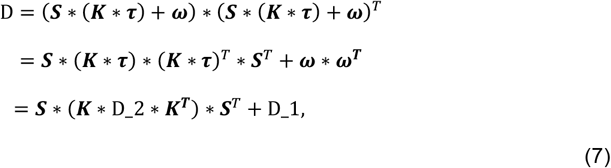

where D_1 = ***ω*** ∗ ***ω***^***T***^ and D_2 = ***τ*** ∗ ***τ***^***T***^ are both diagonal matrices describing the covariance of fluctuations acting on metabolite concentrations and reaction parameters, respectively.

The final equation (7) gives the structure of the fluctuation matrix D in the Lyapunov equation (3), where the matrix D_1 represents the covariance of stochastic fluctuations acting directly on metabolites, and D_2 represents the covariance of stochastic fluctuations in the reaction rate parameters *p*. Assuming that noise acting on the individual components is statistically independent; we can model both as diagonal matrices. Furthermore, enzyme activity or expression data could be used to further constrain the entries of D_2. As shown in Figure 1, we claim that enzymes with large activity variances indicate large fluctuations in relative reaction rate parameters *p*.

### Superpathway based zero and sign structure of the fluctuation matrix

In the previous section, the structure of the fluctuation matrix D is derived for a complete metabolic network. However, in most experiments only a limited number of metabolites is measured which often not provides full coverage of the network. For that situation, we previously developed the COVRECON approach, where we construct a reduced metabolic interaction network for the measured metabolites based on superpathways. Each interaction is a superpathway consists of several reaction steps taking all the possible interaction effects (reactant-product, reactant-reactant, product-product) into account [31, 34]. Notably, each enzyme fluctuation within a superpathway will exert a correlated influence on the metabolites at both ends of the superpathway, introducing off-diagonal components into the structure of the fluctuation matrix D. Consequently, we can map the enzyme-reaction associations to the zero structure of D.

The reaction structure also allows to distinguish positive and negatige off-diagonal elements in D (due to covariance properties, the diagonal elements are all positive). Figure 2 illustrates the three types of superpathways that we need to consider. In the first two types, superpathway X->Y is a one-step pathway. Perturbations on this reaction rate will act positively on the products and negatively on the reactants. Thus, when X and Y are on the same side of the reaction (type 1), the perturbation affects both X and Y in the same way, leading to a positive sign in the element of D corresponding to the interaction between X and Y. Conversely, when X and Y are located on opposite sides of the reaction (type 2), the perturbation influence X and Y differently, leading to a negative sign in the corresponding element. On the other hand, when introducing fluctuation to the metabolite not enzymes, they will enhance the reaction rate and exert a decreasing effect on other metabolites on the same side. Moreover, the influence will pass through the other side of the reaction and influence the metabolites on the other side in the same way. In the more complicated type 3, the superpathway from X to Y consists of several steps of either of the first two types with a number of intermediate metabolites *A*_*i*_. Assuming the enzyme perturbation acts on intermediate step *A*_*k*_ → *A*_*k*+1_, it generates perturbations to both metabolites *A*_*k*_ and *A*_*k*+1_ following the scenarios in type 1 or 2. These perturbations propagate in both directions along the entire superpathway up to X and Y. As illustrated in Figure 2, for the intermediate steps corresponding to type 1, that results in a negative sign; while for intermediate steps of type 2, it results in a positive sign. Consequently, if there are k steps of type 2 where *A*_*i*_ and *A*_*i*+1_ are on the same side of a reaction, the final sign of the D element corresponding to the interaction between X and Y will be −1^*k*+1^.

**Figure 2.**
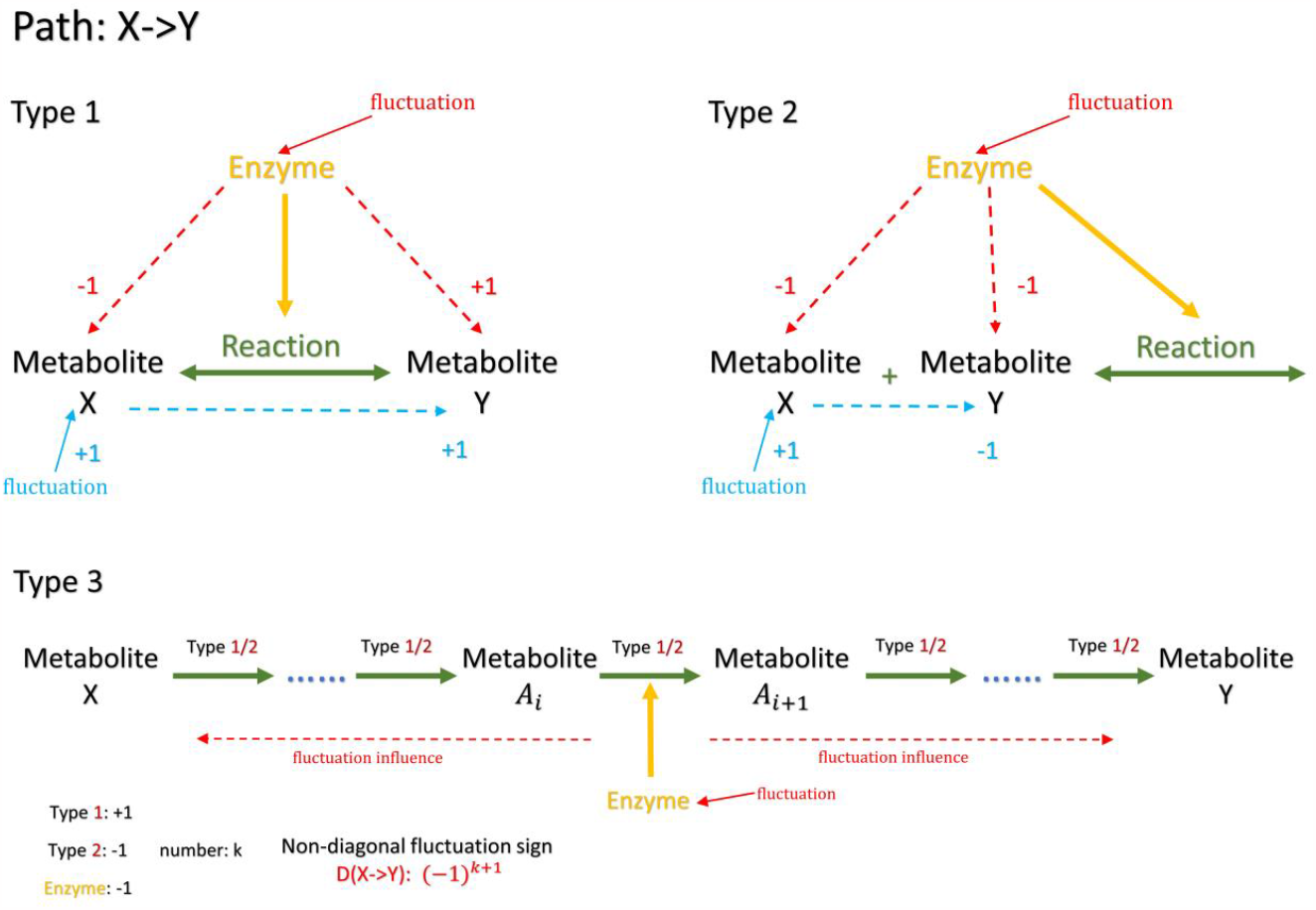
Determine the sign of the non-diagonal D structure in several cases.

The (sign) structure of the fluctuation matrix D is used in the COVRECON Jacobian algorithm to constrain the sampling range for the considered fluctuation values during the inference.

### COVRECON workflow and regression loss based differential Jacobian algorithm

Once we have the structure information of fluctuation matrix D, we follow the same workflow as outlined in COVRECON [31]. It consists of three sub-modules: (i) building of an organism-specific database, (ii) construction of a superpathway-based topological model for metabolite interactions using a pathway search based on the generated organism-specific database, (iii) the regression loss based inverse differential Jacobian computation. Thereby, in step (iii), we sample fluctuation matrices according to the structure of the fluctuation matrix as derived above.

In the inverse Jacobian approach, the Lyapunov Equation (3) is solved using optimization with the data-based covariance matrix C and a sampled fluctuation matrix D. This linear equation can be rewritten in the form

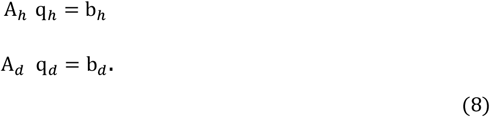

Where A, q and b are computed from corresponding elements from C, J and D respectively. Li, et al. 2023 claimed that under numerical variations in b, coming from sampled fluctuation values, the variation of the regression solution q is much larger compared to the variation in the regression loss *r*. Based on this property, we construct a “regression loss matrix” ***R***^∗^ that aims to capture the relative importance of individual elements in the differential Jacobian, rather than directly calculating the the actual values in the differential Jacobian matrix. Each element of this regression loss matrix ***R***^∗^ is calculated by solving the linear equation (8) with an additional constraint, giving the solution

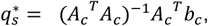

where *A*_*c*_ is constructed by combining *A*_*h*_ and *A*_*d*_ in Equation (8) with the additional constraint that only a single element *J*_*ij*_ may differ between the Jacobians *J*_*h*_ and *J*_*d*_, while all other elements are equal, and *b*_*c*_ = [*b*_*h*_; *b*_*d*_].

The regression loss with respect to that element *J*_*ij*_, corresponding to the element 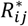 of the regression loss matrix, is then defined as

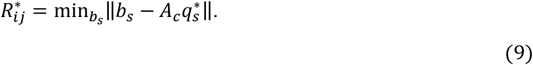

Because we only have the structure information of the fluctuation matrixes *D*_*h*_ and *D*_*d*_, not the actual values, we iterate the regression loss over a number of samples for possible values of the fluctuation matrix that are generated according to its non-zero structure as derived above. The results reported in this paper are based on using 1000 samples. Then, the final 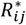 is taken as the minimum regression loss over all samples. In the resulting regression loss matrix ***R***^∗^, larger values then indicate correspondingly larger values in the differential Jacobian matrix. This relation has been shown in [31], where also further details of our inverse differential Jacobian algorithm are described.

### Fluctuation sampling based on enzyme activity data

In the approach described above as well as in the original inverse Jacobian algorithms, the fluctuation matrix D is sampled according to the assumed structure (diagonal or with non-zero off-diagonal elements) with values from an arbitrary “normalized” range between 0 and 1. This assumes that no information on the magnitude of the fluctuations acting on the metabolic network is available. However, in this paper, we take the approach that these fluctuations stem from variability in enzyme activity, for which data can be obtained with omics measurements.

We integrate all the non-diagonal fluctuations inferred from the whole enzyme activity dataset, and sample each non-zero component in *b*_*s*_ with its absolute value between zero and the enzyme activity data related to that component. And the sign of each *b*_*s*_ component is determined as in the Figure 2 or randomly given +1 or -1 if not clearly determined.

### Evaluation of differential Jacobian inference with literature models

To evaluate our algorithm and the potential effect of non-diagonal fluctuations on the differential Jacobian, we consider the four literature models that have also been used in our previous study [31]. Supplementary material S1 gives the model references, and describes how we construct the two conditions required to define a differential Jacobian. And the Jacobian matrixes information are available in Supplementary material S2. The resulting differential Jacobian matrices for these four cases are shown in Figure 3, left column.

**Figure 3.**
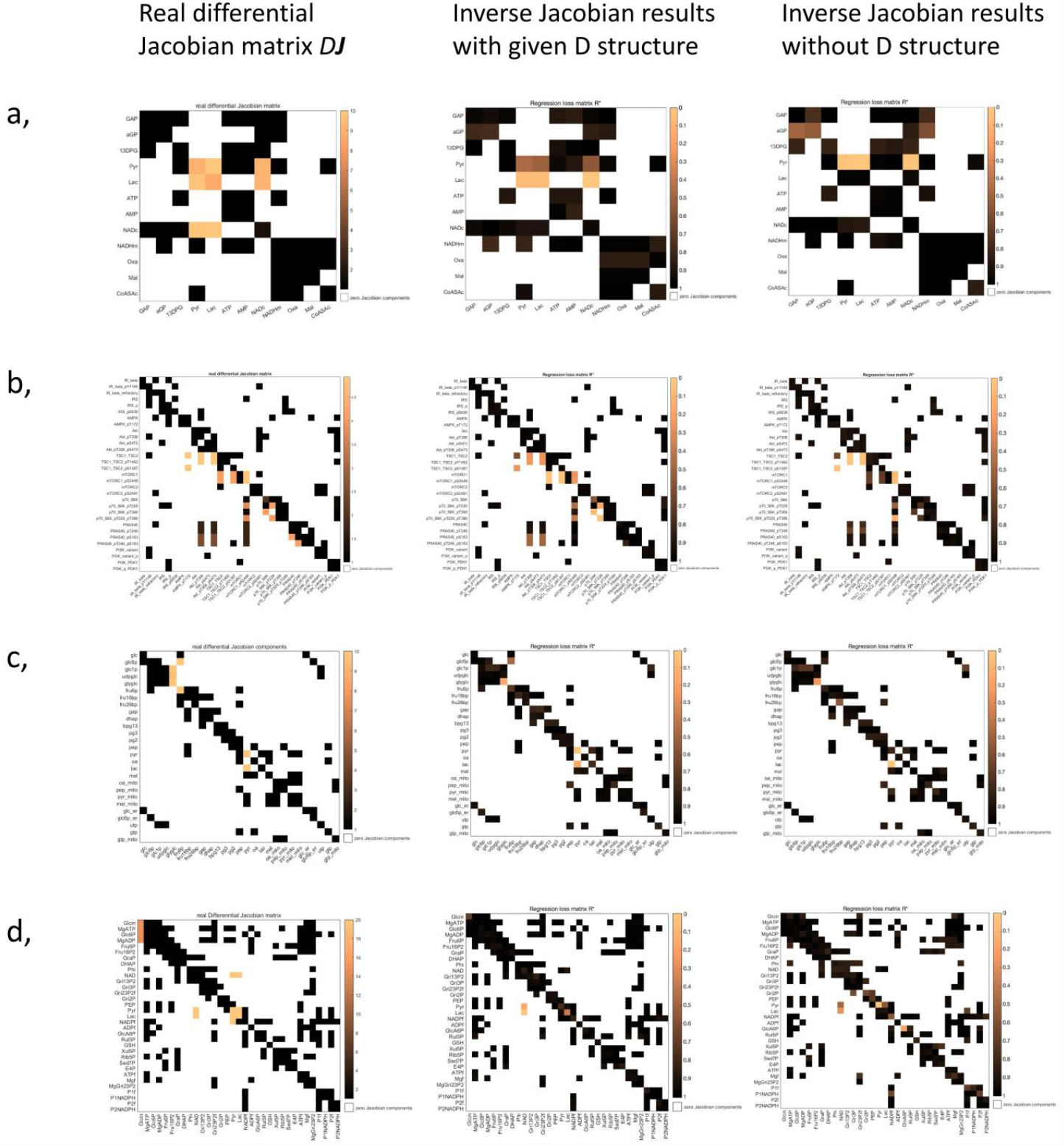
The regression loss Jacobian algorithm evaluation with or without D structure information. The evaluation models are: a, carbohydrate energy metabolism model [50]; b, AMPK-mTOR pathway model [51]; c, hepatic glucose metabolism model [52] and d, blood cell metabolism model [53]. For each subplot a-d, the left is the real differential Jacobian matrix; the middle is the inverse differential Jacobian results with D structure information; and the right is the results without D structure information.

Similar as in previous studies [31, 32, 35, 36], we generate artificial perturbed covariance data for evaluation from the Lyapunov equation through giving random fluctuation matrixes *D*_*h*_ and *D*_*d*_ with different randomness *ε*_*D*_, see [31]. The structure of matrixes *D*_*h*_ and *D*_*h*_ are randomly generated as non-diagonal matrices.

### Inverse differential Jacobian accuracy

We evaluate the accuracy of the inverse differential Jacobian algorithm from three parameters: (1) the magnitude of the fluctuation covariance *ε*_*D*_, (2) the number of off-diagonal components in the fluctuation matrix D, and (3) the fluctuation magnitude of off-diagonal components compared to the diagonal components in fluctuation matrix D, denoted as Md. For each evaluation with specific *ε*_*D*_, N and Md, we calculate the two conditional covariance matrices *C*_*h*_ and *C*_*h*_ from randomly generated non-diagonal fluctuation matrix *D*_*h*_ and *D*_*h*_. Then we conduct the inverse Jacobian analysis with and without taking the structure of the fluctuation matrix D into account. This evaluation process is repeated 200 times. To evaluate how accurately the large components in the real differential Jacobian matrix *D****J*** are identified, we assess the Precision (Positive Predictive Value, PPV) and Recall (True Positive Rate, TPR) of the matrix elements above different threshold values in the regression loss matrix ***R***^∗^. In addition, we assess the precision of the Top 1, Top 3 and Top 5 largest values by rank in ***R***^∗^.

## Results

### Improved differential Jacobian reconstruction when considering off-diagonal fluctuations

As shown in the methods section, if noise acts on reaction parameters, the fluctuation matrix D has a non-diagonal structure. In Equation (6), we derived the resulting structure of D as a sum of two matrices accounting for the perturbations acting on compounds directly (diagonal) and on reaction parameters (both off-diagonal and diagonal), respectively. While the relevant structure information can be determined from just the reaction stoichiometry, determining the relative magnitudes of the off-diagonal elements requires enzyme activity data. In this section, we compare the results of the inverse Jacobian algorithm when taking only the correct structure, but not the magnitude of fluctuations into account, with the previous method were only a diagonal D was used.

For this evaluation, we make use of the four models described in the methods section. For each model, we generate artificial covariance matrix data with perturbations applied to metabolites directly (diagonal fluctuation components) and ¼ of the reactions (off-diagonal components), where these perturbations had the same magnitude. The covariance matrices for the two conditions are computed through the Lyapunov equation with *ε*_*D*_ = 0.4. As shown in Figure 3, the original inverse differential Jacobian algorithm is perturbed by the off-diagonal fluctuations. Nevertheless, we can still achieve good results compared to the real differential Jacobian matrix with the structure information of these off diagonal perturbations.

### Accuracy of the inverse Jacobian algorithm for different fluctuation magnitudes

To evaluate the effect of varying fluctuation magnitudes on the inverse Jacobian algorithm, we generated artifical covariance data for model 1 (carbohydrate energy metabolism) assuming a range of covariance values in the fluctuation matrix D, *ε*_*D*_, ranging from 0.2 to 0.6, and a relative magnitude of off-diagonal fluctuations Md ranging from 0.3 to 2.4. In this analysis, we apply non-diagonal fluctuations to three randomly chosen reactions, and we conduct a second test with six non-diagonal fluctuations applied, which yielded similar results as shown in Supplemental Figure S2. For these data, the inverse Jacobian algorithm has been applied either without any structure information on D, or with assuming the correct structure as described in the methods section. Figure 4 presents the evaluation results, with the first two rows in Figure 4A&4C presenting the inverse Jacobian results without and with D structure respectively, and the third row displaying the difference between the second and the first rows. For each test with specific *ε*_*D*_ and off-diagonal fluctuation magnitude, we repeat the process 200 times and evaluate the large values in the resulting regression loss matrix ***R***^∗^ over the 200 repeats. In Figure 4A, 4B and S1, we compare the precision and recall of correctly identifying differential Jacobian values above a certain threshold, with or without D structure. As an alternative accuracy measure, in Figure 4C, we compare the precision of correctly identifying the top 1, 2 and 5 values in ***R***^∗^.

**Figure 4.**
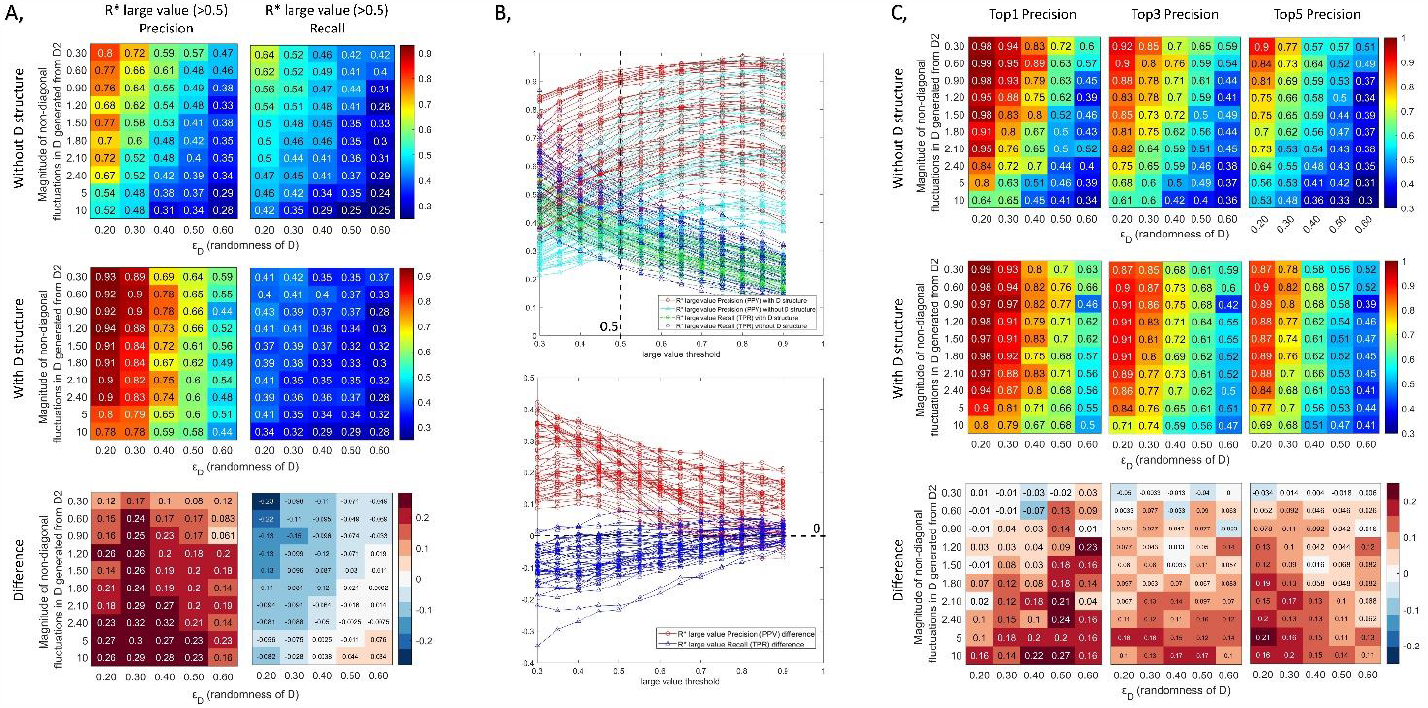
Inverse differential Jacobian algorithm with/without D structure evaluation using various randomness of D *ε*_*D*_ and the magnitude of off-diagonal fluctuations. The evaluation is conducted using the first model in method section with 200 repeats and 3 random enzyme fluctuations applied. A, Precision and Recall of the large values (above 0.5) in ***R***^∗^ over 200 repeats with/without D structure information; B, the line plots of Precision and Recall of the large values in ***R***^∗^ with/without D structure information based on different large value thresholds (0.3-0.9), where the snapshot of value 0.5 refers to A; C, the accuracy of the top 1, top 3 and top 5 large values in ***R***^∗^ over 200 repeats with/without D structure information.

From the results we can conclude that using the structure of the fluctuation matrix D will significantly improve the precision of identifying large values in the differential Jacobian. However, a surprising result is that using the D structure will decrease the recall of the large differential Jacobian values compared to just using a diagonal D. This conclusion remains similar by setting different large value thresholds as shown in Figure 4A (threshold 0.5), S1 (threshold 0.3 & 0.7) and 4B (line plots across thresholds 0.3-0.9). This tells us that using the correct fluctuation matrix structure will sacrifice the hit rate but be able to find the large values in the differential Jacobian with a much higher accuracy. From another point, the sacrifice is worth for a better inverse calculation from the fact that the precision of resulted top 1, 2 and 5 in inverse Jacobian approach is always improved using the D structure information. This is consistent to the results in breast cancer case study, where compared to assuming a diagonal D, the inverse Jacobian results using the D structure will highlight several large values and ignore the others.

Moreover, the results shown in Figure 4A and 4C suggest that the magnitude of covariance values in D has a similar detrimental effect on the inverse differential algorithm in both cases, with the correct structure and with only assuming a diagonal structure. The precision of the results decreases with an increase in the magnitude of non-diagonal fluctuations. However, the decrease is much smaller when using the correct structure of D. Subsequently, the results in the difference plot demonstrate that ignoring the off-diagonal elements in the structure of the fluctuation matrix D generally reduces the reliability of the inverse Jacobian algorithm compared to using the correct structure. However, if the off-diagonal values are smaller than the diagonal values, the results between using the correct structure of D and a diagonal D are comparable.

Finally, using all the four evaluation models, we apply 0.3 magnitude off-diagonal fluctuations to all reactions, and do the inverse Jacobian analysis assuming a diagonal D. Also the evaluation is performed with *ε*_*D*_ = 0.4. As demonstrated in Supplementary Figure S3, without the structure information for D, the new inverse Jacobian algorithm can achieve good results compared to the real differential Jacobian. This reinforces the point that small off-diagonal perturbations can be disregarded in the inverse differential Jacobian approach.

### Accuracy of the inverse Jacobian algorithm with varying number and magnitude of off-diagonal fluctuations

In this section, we analyze the effect of additional assumptions of the fluctuation properties on the accuracy of the Jacobian reconstruction: the magnitude of off-diagonal fluctuations ranging from 0.3 to 10, and the number of off-diagonal perturbations ranging from 6 to 42. This analysis is applied to the first evaluation model. Instead of perturbing reaction parameters, randomly chosen off-diagonal fluctuations are directly added to the fluctuation matrix D. This approach covers all possible non-diagonal fluctuations, not only those from enzyme activities, and can give us a better general understanding of the effect of non-diagonal fluctuations.

Using a similar evaluation as in the previous section, we compare the precision and recall of the large values in the inverse Jacobian over 200 repeats. The results are shown in Figures 5 & S4. As obsered before, using structure information for the fluctuation matirx will increase precision and decrease recall across various magnitudes and numbers of off-diagonal fluctuations compared to inverse Jacobian results assuming only a diagonal structure. Additionally, when assuming a diagonal fluctuation matrix, the inverse Jacobian algorithm accuracy remains relatively consistent across different numbers of small non-diagonal fluctuation, yet, increases with the introduction of more large non-diagonal fluctuations as seen in Figure 5A,s 5C and S4. This is because the non-diagonal components in D originate from the same fluctuations added to several metabolites; consequently, these same fluctuations will also contribute to the related diagonal part of D. Thus, when the number of these fluctuations is large enough, the diagonal part becomes much larger due to the summation of several off-diagonal fluctuations. This further supports our conclusion that assuming a diagonal D remains feasible with only small off-diagonal fluctuations. For an additional validation, we conduct a similar evaluation with *ε*_*D*_ = 0.2 (compared to *ε*_*D*_ = 0.4 in Figure 5), yielding consistent results as presented in Supplemental Figure S5.

**Figure 5.**
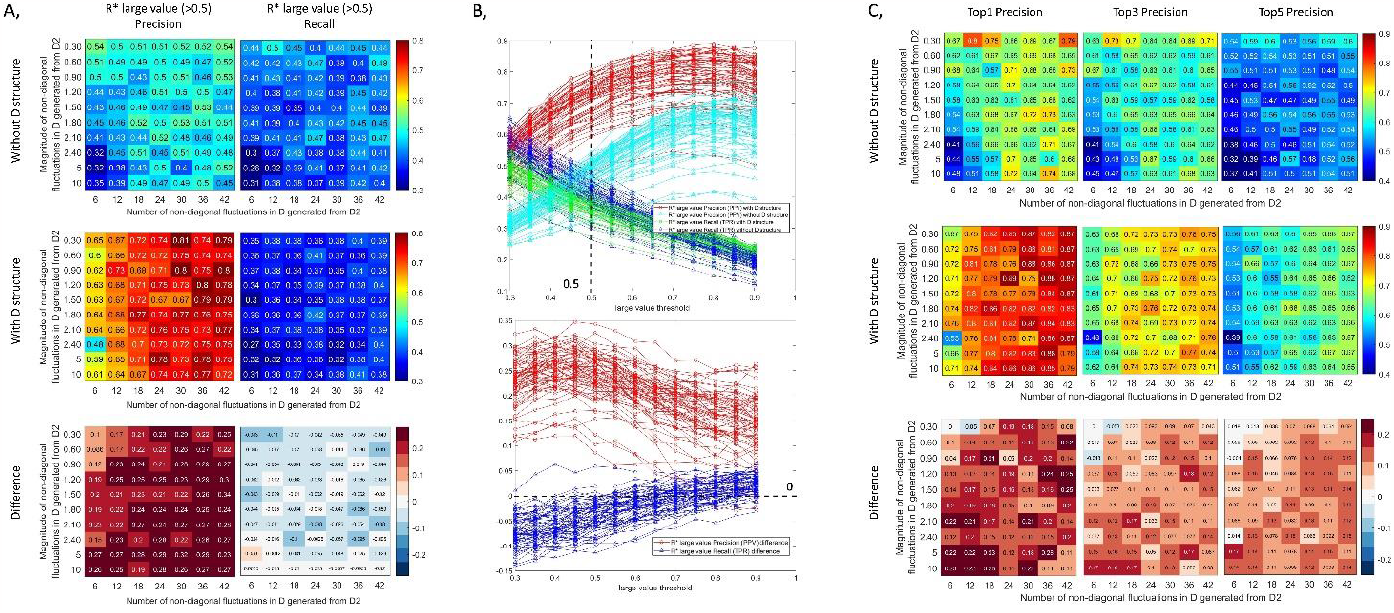
Inverse differential Jacobian algorithm results with/without D structure using a varying number and magnitude of off-diagonal fluctuations. The evaluation is conducted using model 1 form the methods section with 200 repeats and *ε*_*D*_ = 0.4. A, Precision and recall of the large values (above 0.5) in ***R***^∗^ over 200 repeats with/without D structure information; B, the line plots of precision and recall of the large values in ***R***^∗^ with/without D structure information based on different large value thresholds (0.3-0.9), where the snapshot of value 0.5 refers to A; C, the accuracy of the top 1, top 3 and top 5 large values in ***R***^∗^ over 200 repeats with/without D structure information.

### Fluctuation sampling using enzyme activity data further enhances the inverse Jacobian algorithm compared to using a topological D structure

As described in the methods section, enzyme activity data can be used to conduct a more precise sampling of fluctuation values than the purely topology-based sampling used for the previous evaluations. To test the effect of taking this information into account, we vary the number of non-diagonal fluctuations, mixing among fluctuations of small (0.3) and large (2.4) magnitude. Using an approach as in Figure 4, we evaluate the inverse Jacobian algorithm using three sampling strategies: a diagonal D, the topological D structure derived from considering only elements for large fluctuations, and acitivity-based D sampling, which incorporates all fluctuations. Figures 6A, 6C and S6 illustrate that the activity-based D sampling strategy significantly improves the algorithm’s precision compared to using only the topological D structure when the number of large fluctuations is comparatively small. For a comprehensive evaluation, we also conducted this analysis for a wider spread of fluctuation magnitudes with 10 vs. 0.3, and *ε*_*D*_ values of 0.2 and 0.4, respectively. Supplementary Figures S7 & S8 present the results for these scenarios, which are in line with the results shown in Figure 6.

**Figure 6.**
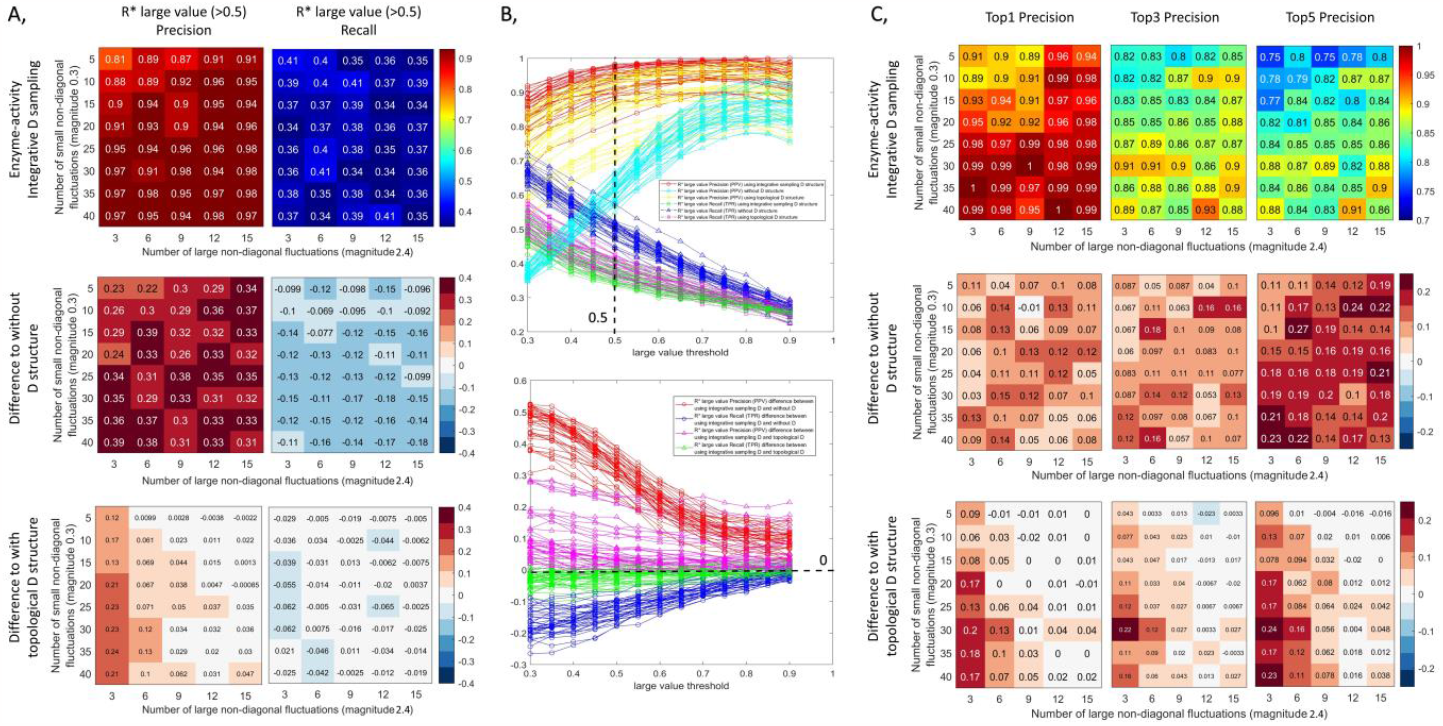
Inverse differential Jacobian results using various number of large (magnitude: 2.4) and small (magnitude: 0.3) non-diagonal fluctuations from three fluctuation matrix assumptions: enzyme-activity based D sampling, topological D and diagonal D. The evaluation is conducted using model 1 from the methods section with 200 repeats and *ε*_*D*_ = 0.2. A, Precision and recall of the large values (above 0.5) in ***R***^∗^ over 200 repeats with/without D structure information; B, the line plots of Precision and Recall of the large values in ***R***^∗^ with/without D structure information based on different large value thresholds (0.3-0.9), where the snapshot of value 0.5 refers to A; C, the accuracy of the top 1, top 3 and top 5 large values in ***R***^∗^ over 200 repeats with/without D structure information.

### Application to a breast cancer dataset

Finally, we apply the inverse Jacobian approach to a breast cancer dataset that covers two cell lines [46]: a non-tumorigenic breast epithelial cell line (MCF102A) and a pleural effusion metastasis of a breast adenocarcinoma (MCF7). The objective is to find large values in the differential Jacobian between these cell lines from metabolomics covariance data. In addition to metabolites, the authors also measured transcriptomics data, and provided a genome-scale metabolic network model (RECON3D_301_hgnc_id.xml) in which the reactions are annotated with gene IDs corresponding to the identifiers used in transcriptomics. Using the Cobra toolbox [54], we are able to generate from the transcriptomics data a value for each reaction to represent the enzyme activity for that reaction. Even though transcript levels are not fully representative of enzyme activity, we use this value here as an upper bound on enzyme activity fluctuations during the sampling. Collecting the enzyme activity profiles for the entire genome-scale model, we can then compute the variance of each enzyme activity separately for the two different cell lines. The histograms illustrating the variance in enzyme activity values for the two cell lines can be found in Figure S9. Notably, there are a significant portion of small-perturbed enzymes while only a very limited number of enzymes exhibit activity variances exceeding 200.

Using the default setting in COVRECON toolbox, we first construct the metabolic interaction network for the metabolomics dataset, where the interaction information can be checked in Supplementary material S2. Consequently, we map the calculated variances of the enzyme activities to the determined metabolic interactions and construct the structure of the fluctuation matrix D as shown in Figure 7. We perform the inverse differential Jacobian algorithm with the enzyme-activitiy based D sampling strategy. We compare different scalings of the fluctuations vs. each other by setting the magnitude of fluctuations affecting metabolites directly (D_1) equal to fluctuations from enzyme activity (D_2) with variance values equal to 500, 200, and 10, respectively. The results are then compared to the results obtained in our previous study using a diagonal D [31], as depicted in Figure 8.

**Figure 7.**
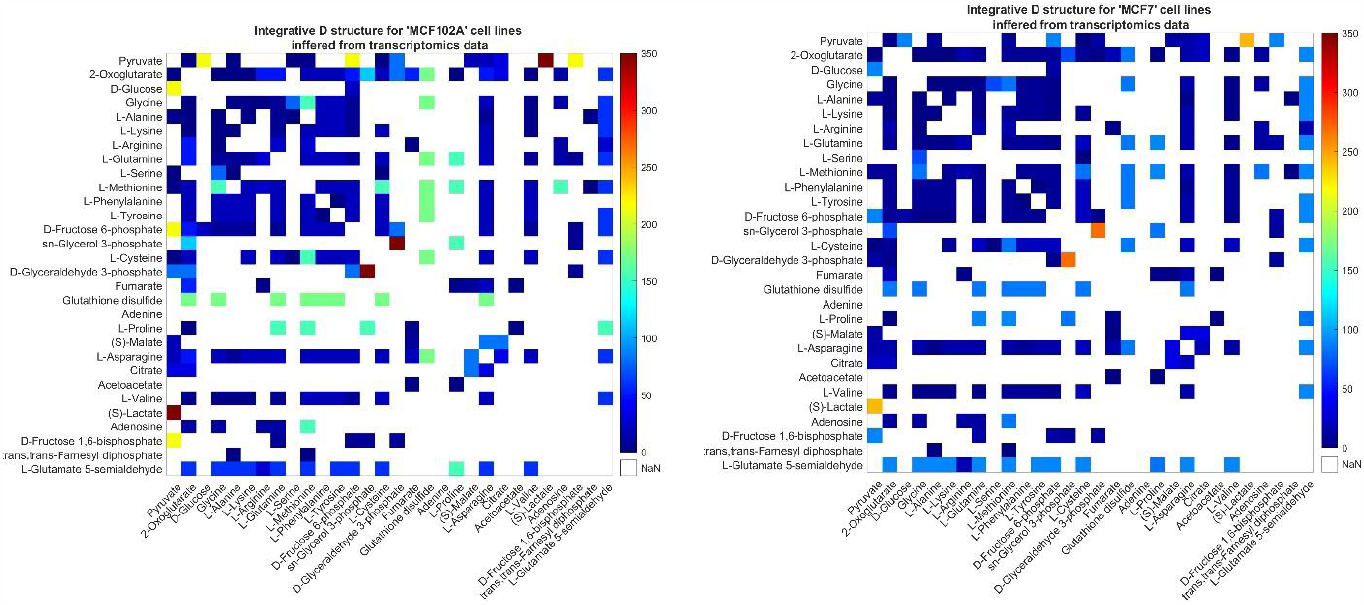
The inferred fluctuation matrix structure for the two different cell lines. (left: D_h_ for MCF 102A and right: D_d_ for MCF 7).

**Figure 8.**
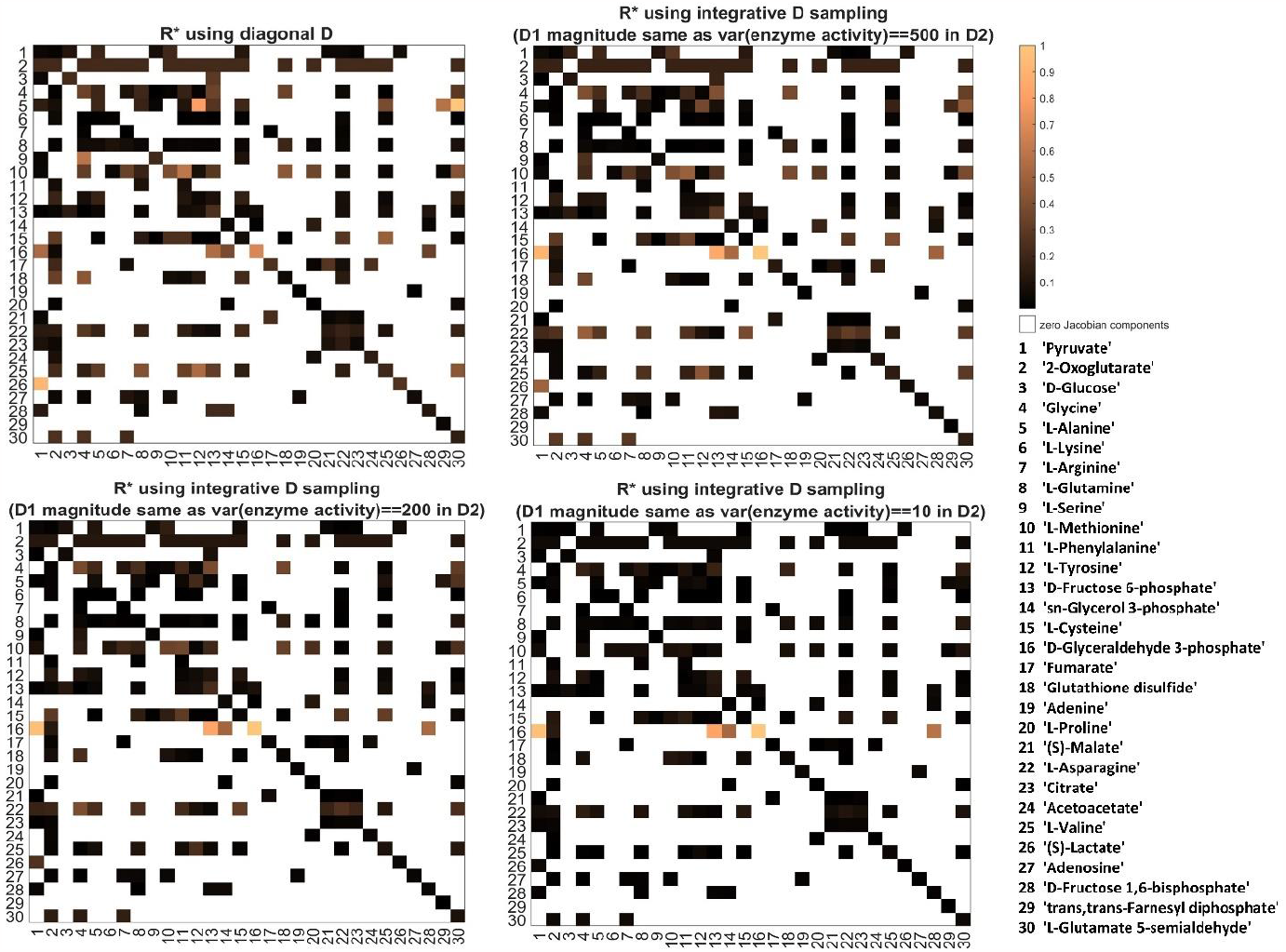
Inverse differential Jacobian analysis on the breast cancer dataset. The four panels show inference results using a diagonal D, and three sets of enzyme-activity based D_h_ and D_d_ sampling, where the magnitude of diagonal fluctuations in D1 is scaled equal to fluctuations from enzyme activities with variances of 500, 200, and 10, respectively.

It is apparent that when scaling the magnitude of diagonal fluctuations D1 to be the same as a large non-diagonal enzyme activity fluctuation in D2, with variance of 500, the inverse Jacobian algorithm result with our new approach is close to the original inverse Jacobian algorithm result using a diagonal D sampling, as shown in right top subplot in Figure 8. This result assumes that metabolite fluctuations play an important role. Conversely, when scaling the magnitude of diagonal fluctuations D1 magnitude equal to a small non-diagonal enzyme activity fluctuation with variance of 10, the results from the two approaches differ significantly. In this scenario, enzymes fluctuations play the dominant role. In line with the reduced recall seen in the previous evaluations, fewer, but more pronounced metabolite interactions are inferred to take a large value in the differential Jacobian. Following the increased precision of the enzyme-activity based inference seen in the previous tests, we postulate that these elements more reliably correspond to an actual change in metabolic interactions between the considered cell lines, even if we cannot determine the actual ratio between fluctuations acting directly on metabolites and those acting on enzyme activities. The inverse Jacobian results show that the main dynamic difference between the cell lines lies on the metabolite D-Glyceraldehyde 3-phosphate, which is also identified as a biomarker in their original t-test [46]. Several interactions are identified as differences between the two cell lines where relevant enzymes activity also show significant difference [31]. The circular plots where one can check the superpathway information interactively are shown in Supplementary Figure S10 and available as matlab figure format in Supplementary material S2.

Interestingly, the two largest enzyme fluctuations in the super-pathway interactions here considered are also identified as large values in the inverse differential Jacobian result: sn-Glycerol 3-phosphate->D-Glyceraldehyde 3-phosphate and Pyruvate->(S)-Lactate. This suggests that indeed the key fluctuations in metabolic variance observed between the cell lines originate from enzyme activity fluctuations.

## Conclusions

In this article, we have introduced a novel inverse differential Jacobian approach that leverages more information about the structure of the fluctuation matrix, including reaction network structure and enzyme activity data. A fluctuation sampling strategy is applied that makes use of suitable data for estimating variance in enzyme activity. We assessed the influence of these non-diagonal fluctuations on the inverse Jacobian algorithm based on three key factors: (1) the magnitude of covariance values, (2) the number of non-zero off-diagonal components in the fluctuation matrix D, and (3) the magnitude of off-diagonal components compared to that of diagonal components. The main findings are:

1. While sacrificing the positive hit rate, incorporating the non-diagonal D structure into the analysis significantly enhances the precision of the inverse Jacobian algorithm. Using an enzyme-activity based sampling strategy further improves the algorithm’s accuracy compared to using only a topological structure of the fluctuation matrix.
2. There are two cases in which assuming a diagonal fluctuation structure remains feasible for the inverse differential Jacobian analysis: (1) When non-diagonal fluctuations are relatively small in comparison to direct metabolite fluctuations; (2) when there are numerous non-diagonal fluctuations with similar magnitudes.

This approach integrates the different OMICS datasets into the inverse Jacobian analysis, contributes to our deeper understanding of metabolic network interaction and dynamics, and enhances the robustness of the inverse Jacobian inference.

## Data availability

The data underlying this article are available in its online supplementary material. And the original data of breast cancer case study can be accessed in the reference [46]. The matlab code is available in https://bitbucket.org/mosys-univie/covrecon/.

## Supplementary Material S1

### Evaluation literature models

To evaluate our algorithm and the potential effect of non-diagonal fluctuations on the differential Jacobian, we consider the four literature models that have also been used in our previous study [1]. We describe how to generate the two conditional Jacobians as following:

1. A model of carbohydrate energy metabolism [2]: One condition is obtained from the nominal parameter values; for the other condition we increased the reaction rate parameter in R2, Pyrute+NADHc→Lactose+NADc, five-fold.
2. AMPK-mTOR pathway model [3]: The paper describes a wild type and an mTOR knockout model based on time-series experimental data; these two variants are used to define the differential Jacobian.
3. Hepatic glucose metabolism model [4]. The first condition is from nominal parameters. We applied a two-fold parameter change for the reaction rate of the second reaction R2 for the second condition.
4. Large-scale blood cell metabolism model [5]. We introduced a five-fold increase to several components of the Jacobian matrix directly.

These Jacobian matrixes are available in Supplementary material S2.

**Figure S1.**
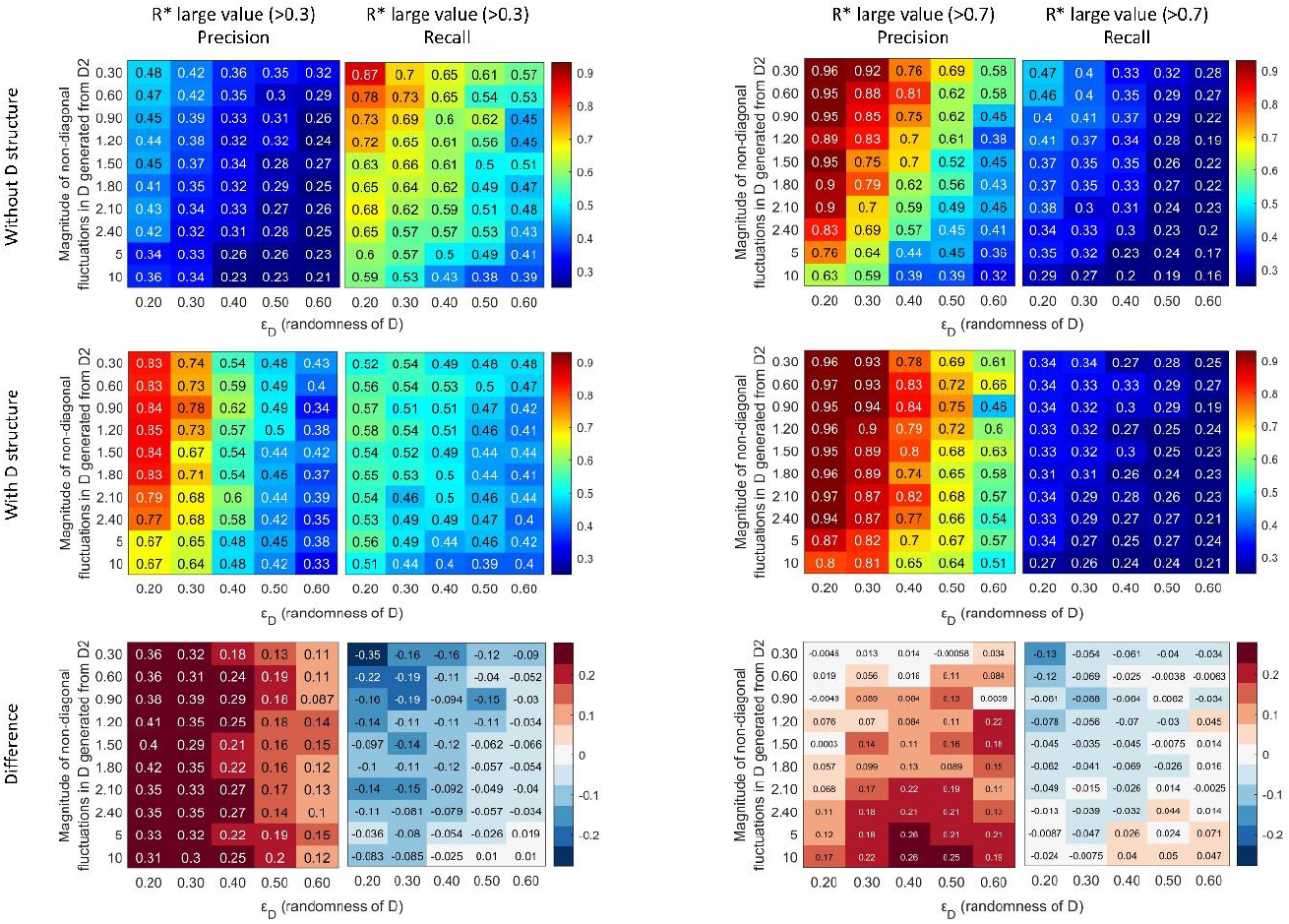
Precision and Recall of the large values in *R*^∗^ based on different thresholds (left: 0.3, right: 0.7). Color code refers to Figure 4A.

**Figure S2.**
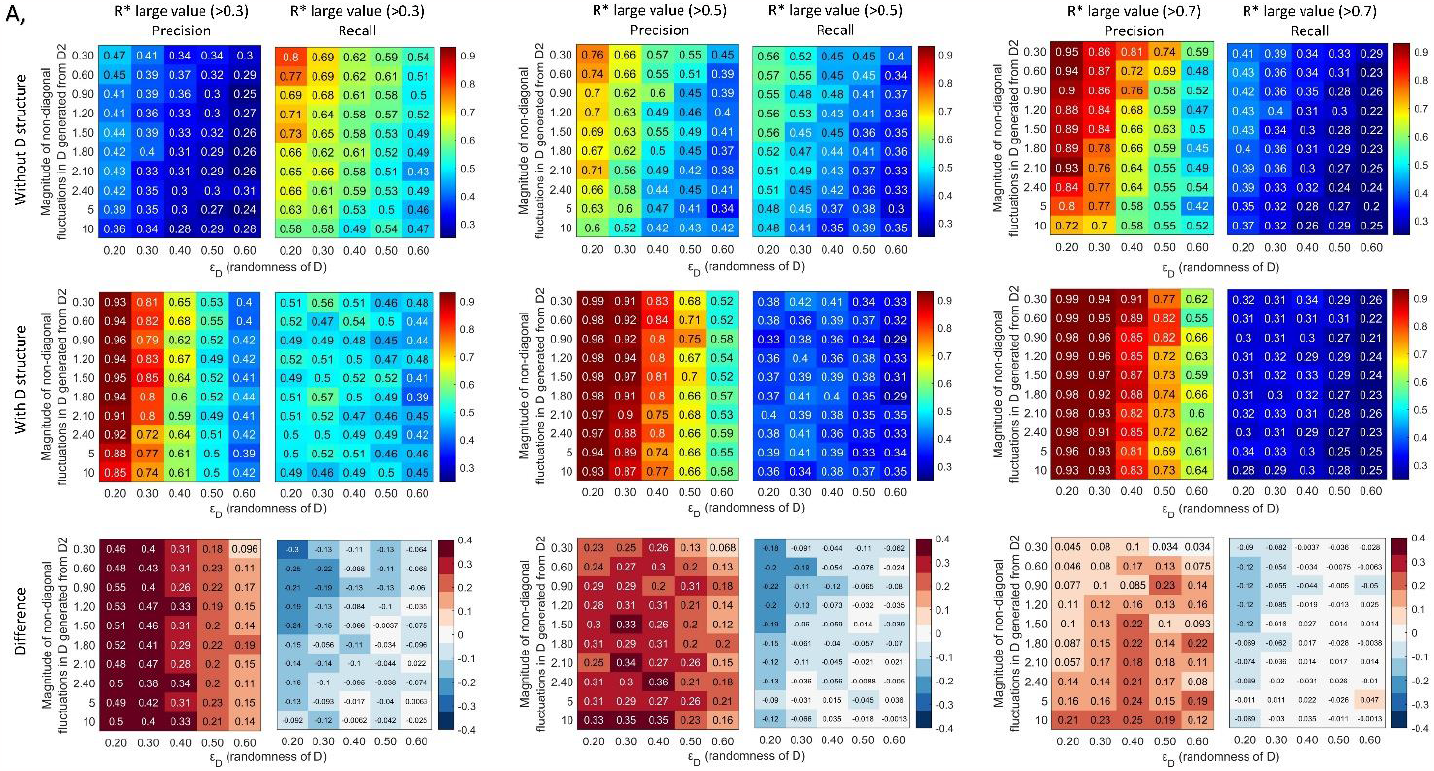
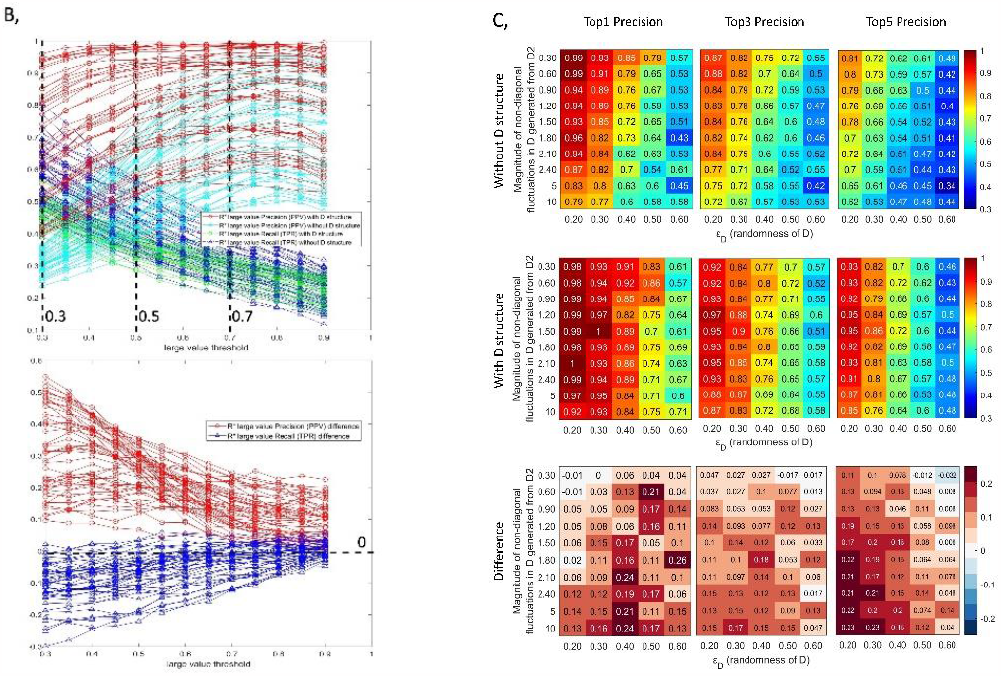
Inverse differential Jacobian algorithm with/without D structure evaluation using various randomness of D *ε*_*D*_ and the magnitude of off-diagonal fluctuations. The evaluation is conducted using the first model in method section with 200 repeats and 6 random enzyme fluctuations applied. A, Precision and Recall of the large values (above 0.5) in ***R***^∗^ over 200 repeats with/without D structure information; B, the line plots of Precision and Recall of the large values in ***R***^∗^ with/without D structure information based on different large value thresholds (0.3-0.9), where the snapshots of value 0.3, 0.5 and 0.7 refer to A; C, the accuracy of the top 1, top 3 and top 5 large values in ***R***^∗^ over 200 repeats with/without D structure information.

**Figure S3.**
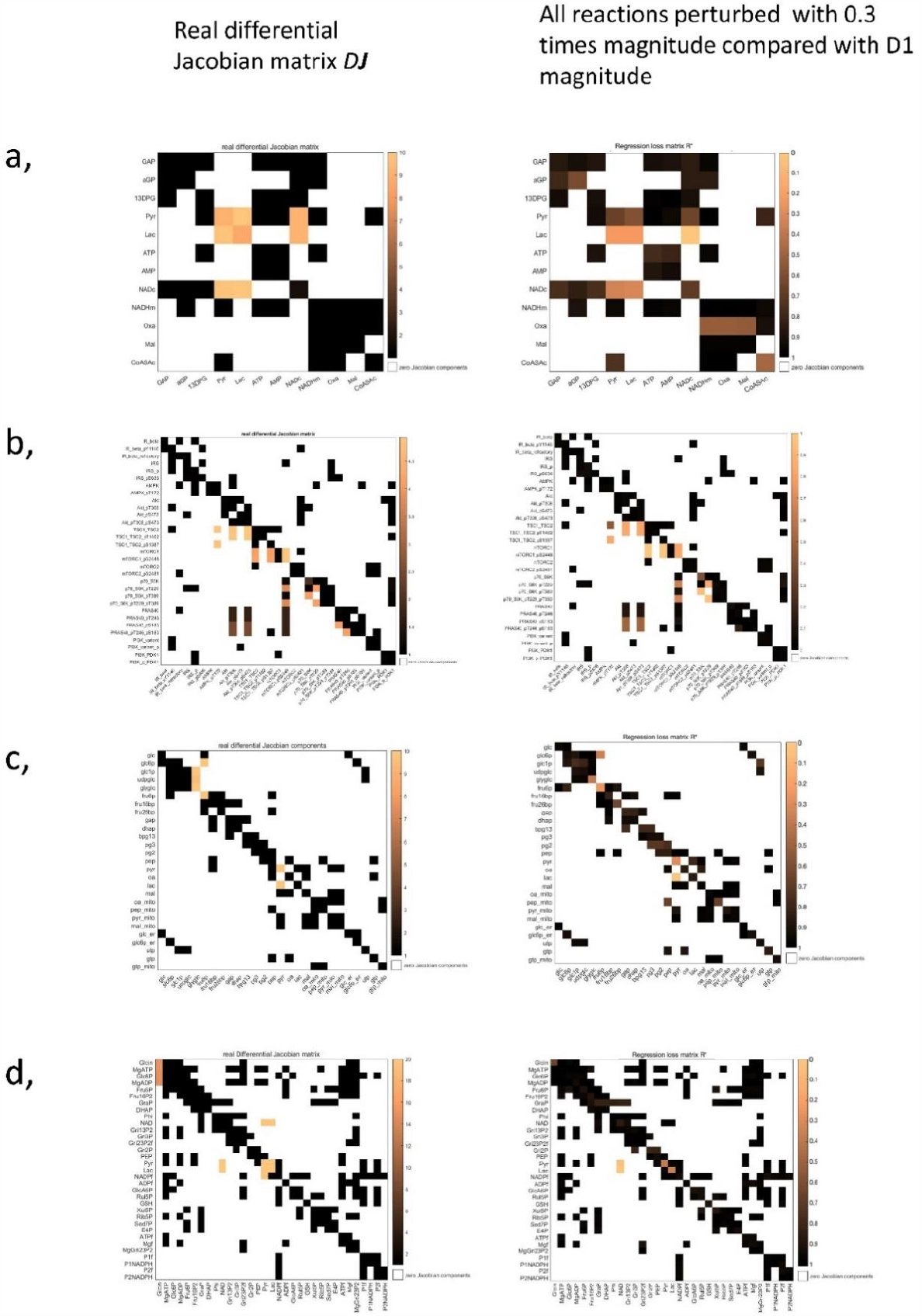
The regression loss Jacobian algorithm evaluation assuming a diagonal fluctuation matrix D with *ε*_*D*_ = 0. 4. For each test, all reactions are perturbed using a 0.3 magnitude fluctuation compared the metabolites fluctuation.

**Figure S4.**
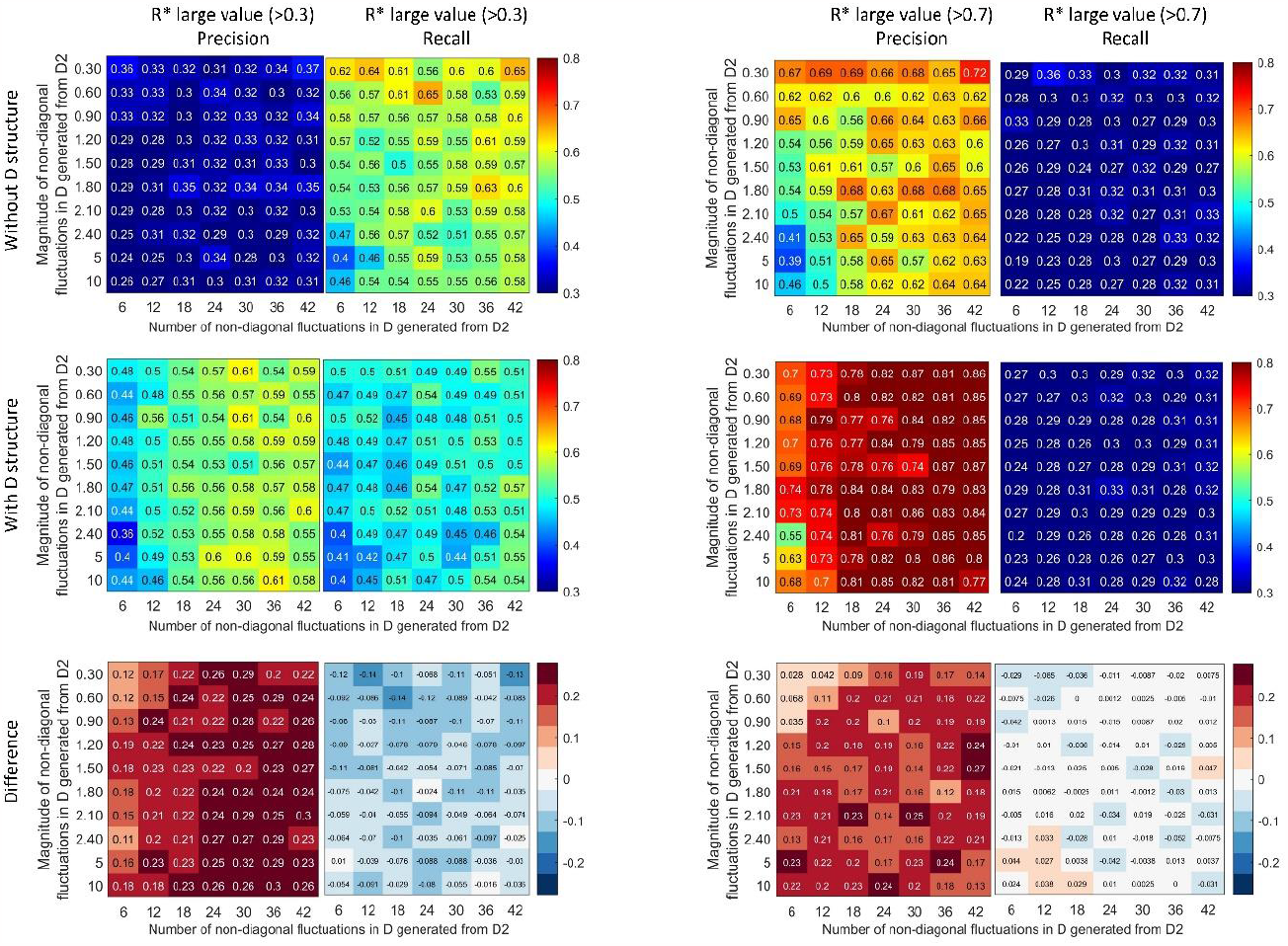
Precision and Recall of the large values in *R*^∗^ based on different thresholds (left: 0.3, right: 0.7). Color code refers to Figure 5A.

**Figure S5.**
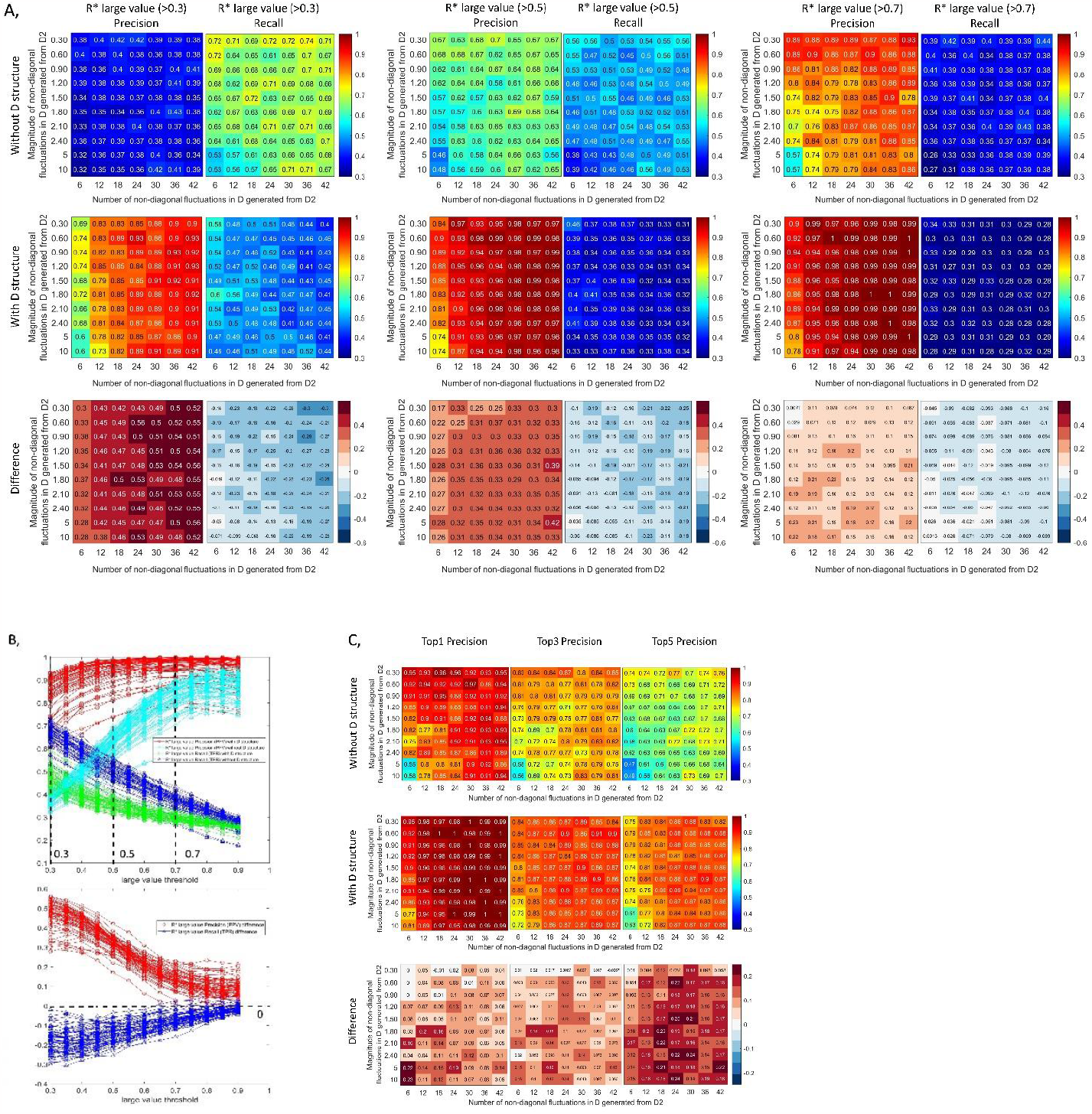
Inverse differential Jacobian algorithm with/without D structure evaluation using various number and magnitude of off-diagonal fluctuations. The evaluation is conducted using the first model in method section with 200 repeats and *ε*_*D*_ = 0.2. A, Precision and Recall of the large values (above 0.5) in ***R***^∗^ over 200 repeats with/without D structure information; B, the line plots of Precision and Recall of the large values in ***R***^∗^ with/without D structure information based on different large value thresholds (0.3-0.9), where the snapshot of value 0.5 refers to A; C, the accuracy of the top 1, top 3 and top 5 large values in ***R***^∗^ over 200 repeats with/without D structure information.

**Figure S6.**
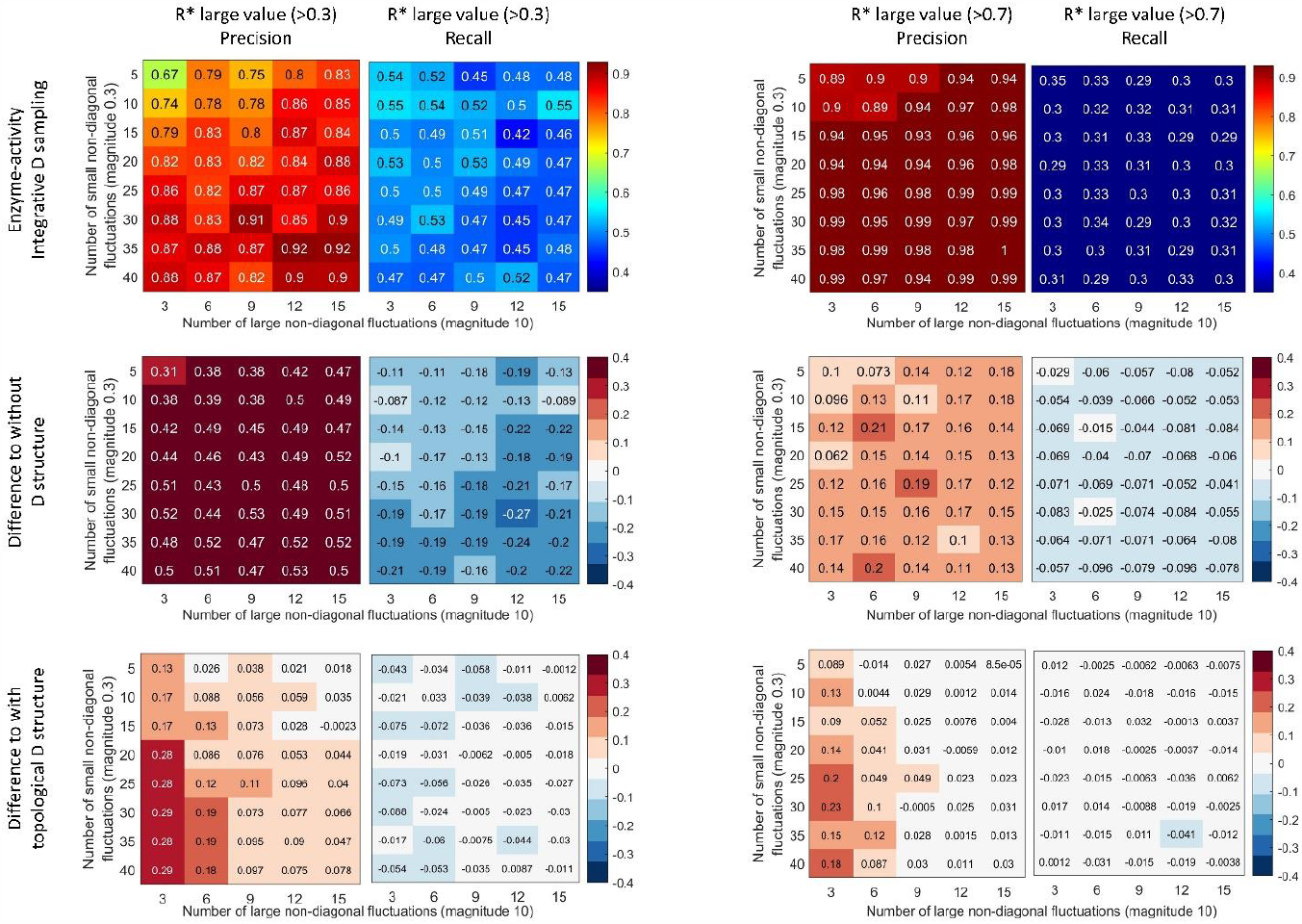
Precision and Recall of the large values in *R*^∗^ based on different thresholds (left: 0.3, right: 0.7). Color code refers to Figure 6A.

**Figure S7.**
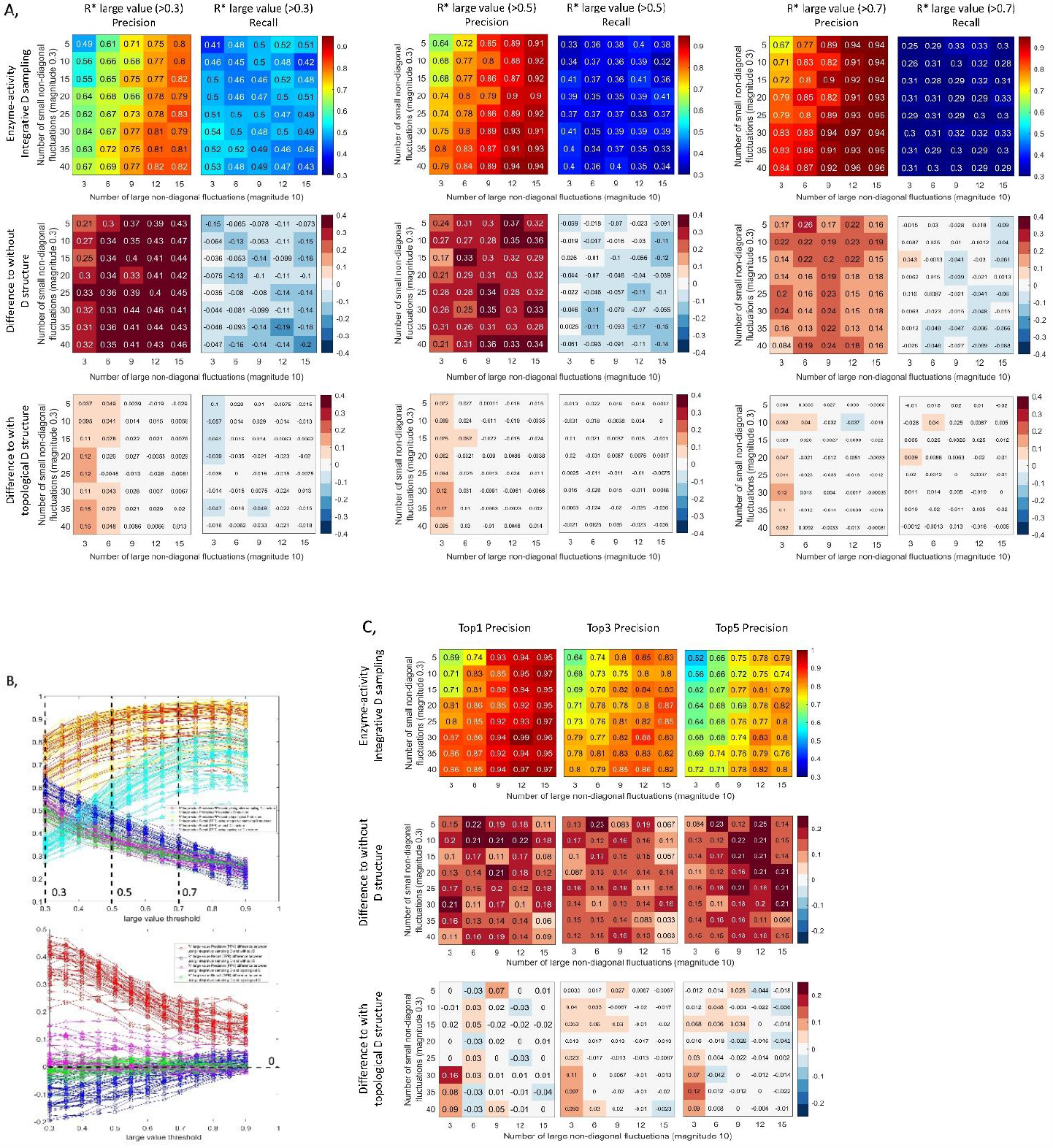
Inverse differential Jacobian algorithm evaluation using various number of large (magnitude: 10) and small (magnitude: 0.3) non-diagonal fluctuations using three strategies: integrative D sampling, topological D and without D structure. The evaluation is conducted using the first model in method section with 200 repeats and *ε*_*D*_ = 0.2. A, Precision and Recall of the large values (above 0.5) in ***R***^∗^ over 200 repeats with/without D structure information; B, the line plots of Precision and Recall of the large values in ***R***^∗^ with/without D structure information based on different large value thresholds (0.3-0.9), where the snapshot of value 0.5 refers to A; C, the accuracy of the top 1, top 3 and top 5 large values in ***R***^∗^ over 200 repeats with/without D structure information.

**Figure S8.**
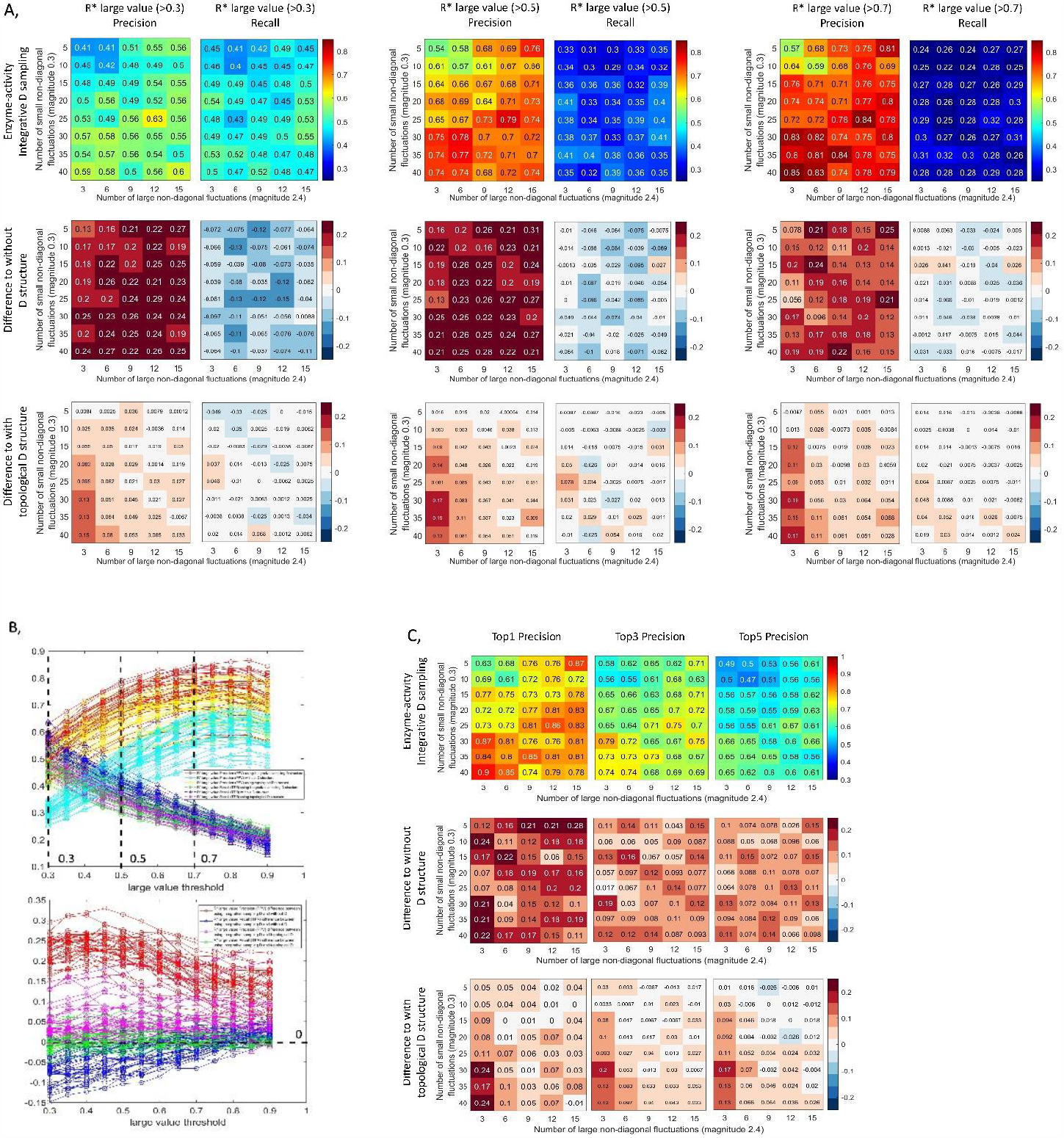
Inverse differential Jacobian algorithm evaluation using various number of large (magnitude: 2.4) and small (magnitude: 0.3) non-diagonal fluctuations using three strategies: integrative D sampling, topological D and without D structure. The evaluation is conducted using the first model in method section with 200 repeats and *ε*_*D*_ = 0.4. A, Precision and Recall of the large values (above 0.5) in ***R***^∗^ over 200 repeats with/without D structure information; B, the line plots of Precision and Recall of the large values in ***R***^∗^ with/without D structure information based on different large value thresholds (0.3-0.9), where the snapshot of value 0.5 refers to A; C, the accuracy of the top 1, top 3 and top 5 large values in ***R***^∗^ over 200 repeats with/without D structure information.

**Figure S9.**
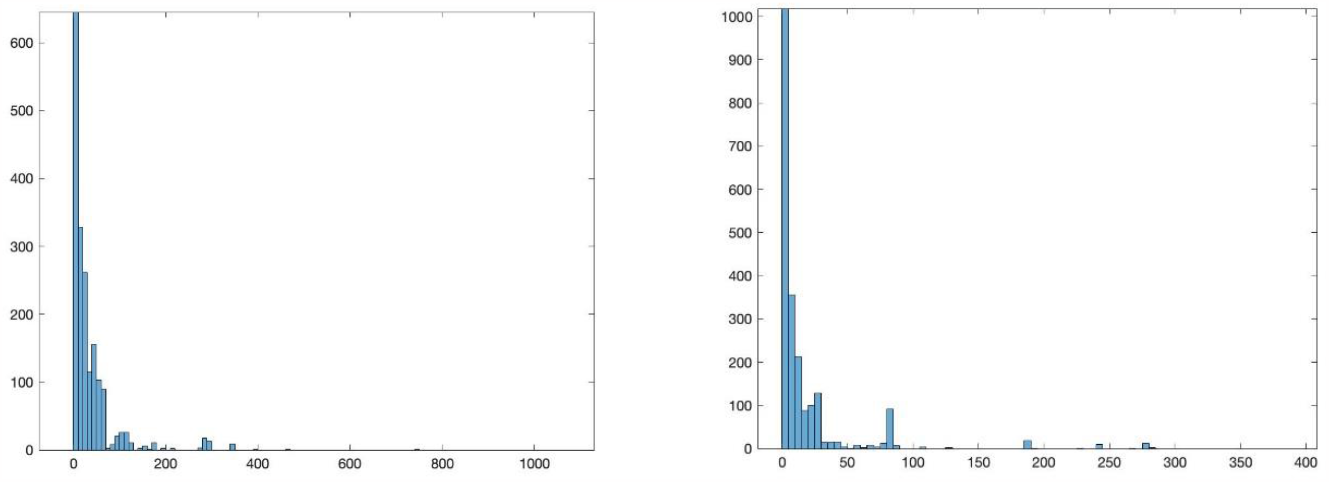
The histogram plots of the activities variances of the two cell lines enzymes.

**Figure S10.**
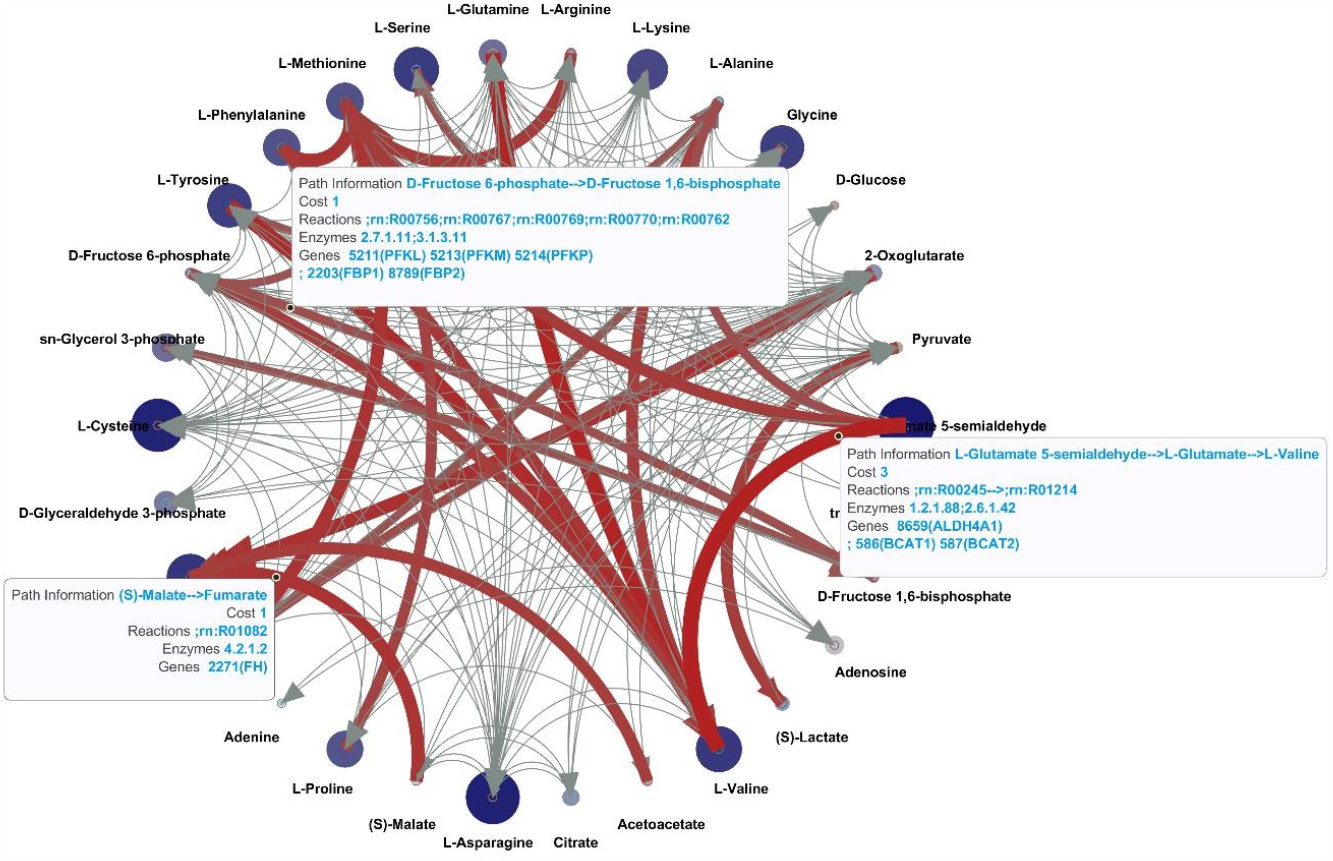
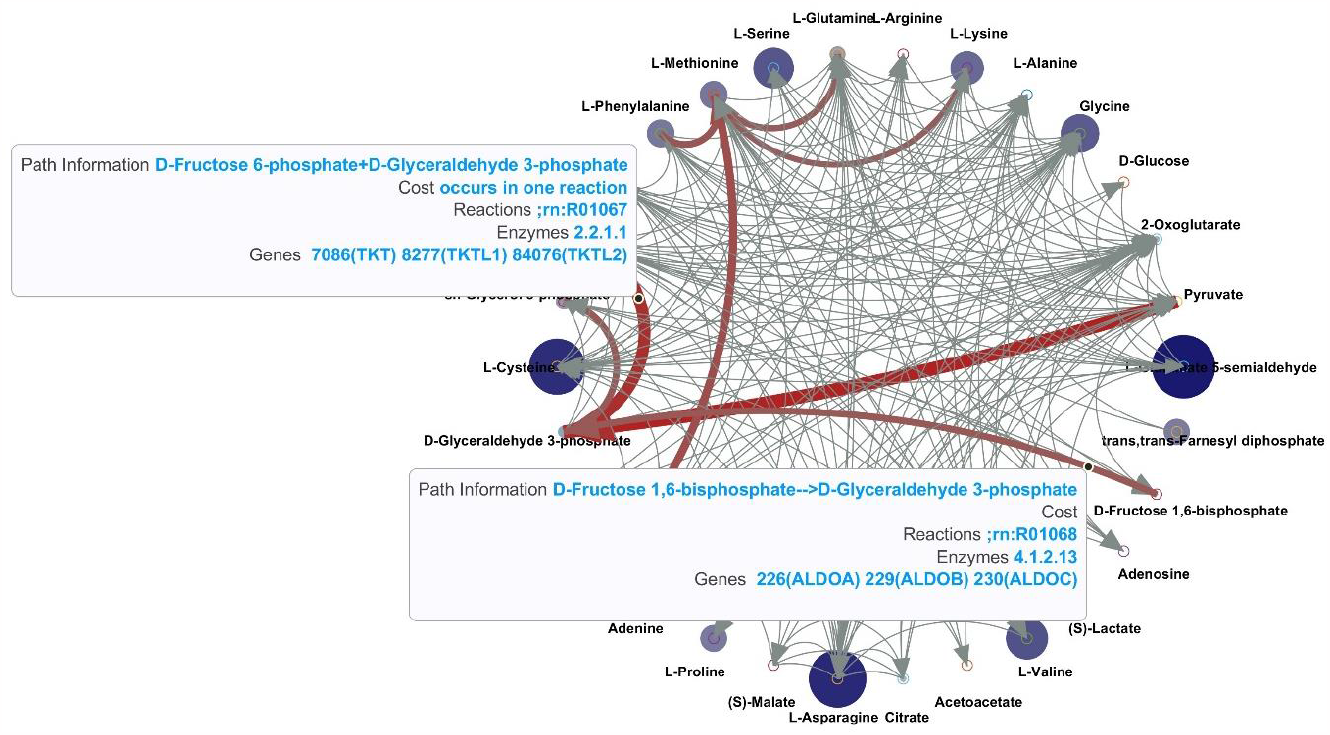
The interactive plots of breast cancer inverse Jacobian analysis. Top plot uses a diagonal D assumption and bottom plot uses enzyme-activity integrative D sampling. The circular plot can be interactively checked by clicking the interactions (lines) through the matlab figure format result available in Supplementary material S2.

