## Supplementary_material_S2 for "Enzyme fluctuations data improve inference of metabolic interaction networks with an inverse differential Jacobian approach": Enzyme_Fluctuation_Inverse_Jacobian_algorithm.m

```
function R=Enzyme_Fluctuation_Inverse_Jacobian_algorithm(C1,C2,J_struc,name_v,E_1,E_2)
% C1 C2 are two covariance matrices of the system at two conditions;
% J_struc is the Jacobian matrix structure information; 
% name_v is the names of all metabolites measurements;
% E_1,E_2 are the enzyme fluctuation matrices of the two conditions


warning('off')
%  test  
J=J_struc;
J1=J;
J2=J;

Jac1=J1;
Jac2=J2;


  
n=length(Jac1);
J=ones(n);
J(Jac1==0)=0;


D=eye(n);
D_t1=zeros(n*(n+1),1);
D_t1(1)=-1;
D_t1(1+n*(n+1)/2)=-1;
t=1;
for i=1:(n-1)
    t=t+n+1-i;
    D_t1(t)=-1;
    D_t1(t+n*(n+1)/2)=-1;
end
       
D1=eye(n);
[A1,B1]=materix_trans(C1,D1,Jac1);


J1=A1\B1';


Jac1=J_struc;

n=length(Jac1);

[A1,B1]=materix_trans(C1,D1,Jac1);


ss_min=[];
ss_min1=[];
ss_min2=[];

 s_size=1000; 
 rand_D=[];
 

D_t1=zeros(n*(n+1),1);
D_t1(1)=-1;
D_t1(1+n*(n+1)/2)=-1;
t=1;
for i=1:(n-1)
    t=t+n+1-i;
    D_t1(t)=-1;
    D_t1(t+n*(n+1)/2)=-1;
end


for uu=1:s_size
            
            aa=0.1+rand(n,n);
            aa=(aa+aa')/2;
            bb=0.1+rand(n,n);
            bb=(bb+bb')/2;
            D_samp=E_1;
            D_samp(eye(n)==1)=0;
            
            D_samp1=D_samp.*aa;
            for i=1:n
                D_samp1(i,i)=sum(abs(D_samp1(i,:)))+rand();
            end

            D_samp=E_2;
            D_samp(eye(n)==1)=0;
            D_samp2=D_samp.*bb;
            for i=1:n
                D_samp2(i,i)=sum(abs(D_samp2(i,:)))+rand();
            end
kk=1;
 D_t1=[];
                D_t2=[];

            for i=1:n
    for j=i:n
        D_t1(kk)=D_samp1(i,j);
        kk=kk+1;
    end
            end    

            kk=1;
            for i=1:n
    for j=i:n
        D_t2(kk)=D_samp2(i,j);
        kk=kk+1;
    end
            end 
            D_t=-[D_t1,D_t2];
            D_t=D_t';
            
            D_t=D_t/sqrt(mean(D_t.^2));
            rand2_D{uu}=D_t;      
end


    [A1,B1]=materix_trans(C1,D1,Jac1);

J1=A1\B1';

b=msgbox('Calculating Inverse Jacobian Regression Loss R*...');

parfor i=1:length(J1)
    warning('off')
    
   D2=D1;
    [A1,B1]=materix_trans(C1,D1,Jac1);
    [A2,B2]=materix_trans(C2,D2,Jac1);

       A_total=[];
       group_now=[];
for j=1:length(J1) group_now(j)=j; end
group_now(i)=[];
       A_diff=[A1(:,group_now),zeros(n*(n+1)/2,length(group_now));zeros(n*(n+1)/2,length(group_now)),A2(:,group_now)];
       A1(:,group_now)=[];
       A2(:,group_now)=[];
       A1_p=A1;
        A2_p=A2;
        A_total=[A_diff,[A1_p;A2_p]];
        
        ss_all=[];
          for j=1:s_size
            D_t=rand_D{j};

            [b,bint,r,rint,stats] = regress(D_t,A_total);
            
           ss_all(j)= stats(4);
            
          end
          ss_all1=ss_all;
            ss_all=sort(ss_all);
             ss_min(i)=mean(ss_all(1:1)) ;
        %     find(ss_all1==ss_min(i))
            ss_min1(i)=mean(ss_all(1:10)) ;
            ss_min2(i)=mean(ss_all(1:20)) ;

J_total=A_total\D_t;

end
delete(b)
KK=[];
for i=1:length(J1) KK(i)=i; end
SS1=[KK;ss_min]';
SS1=sortrows(SS1,2,'descend'); 
SS2=[KK;ss_min1]';
SS2=sortrows(SS2,2,'descend'); 
SS3=[KK;ss_min2]';
SS3=sortrows(SS3,2,'descend'); 
SS_0_6=[SS1,SS2,SS3];


xvalues=name_v;
yvalues=name_v;
Jac1=J_struc;
index=find(Jac1'~=0);


max1=max(SS_0_6(:,6));
min1=min(SS_0_6(:,6));

J_cal=zeros(length(Jac1));
for i=1:length(index)
    a=(SS_0_6(i,6)-min1)/(max1-min1); 
J_cal(index(SS_0_6(i,5)))=a;
end


max1=max(SS_0_6(:,4));
min1=min(SS_0_6(:,4));

J_cal1=zeros(length(Jac1));
for i=1:length(index)
    a=(SS_0_6(i,4)-min1)/(max1-min1); 
J_cal1(index(SS_0_6(i,3)))=a;
end

max1=max(SS_0_6(:,2));
min1=min(SS_0_6(:,2));

J_cal2=zeros(length(Jac1));
for i=1:length(index)
    a=(SS_0_6(i,2)-min1)/(max1-min1); 
J_cal2(index(SS_0_6(i,1)))=a;
end

J_cal=(J_cal+J_cal1+J_cal2)/3;
J_cal((J_cal==0))=NaN;
% figure(1)
% h1=heatmap(xvalues,yvalues,J_cal');


R=J_cal';


end


function loss_r1=loss_r(D_dia,A_total)
warning('off')
D_dia=D_dia/sum(D_dia)*12;
D_t2=zeros(42,1);
D_t2(1)=D_dia(1);
D_t2(7)=D_dia(2);
D_t2(12)=D_dia(3);
D_t2(16)=D_dia(4);
D_t2(19)=D_dia(5);
D_t2(21)=D_dia(6);
D_t2(22)=D_dia(7);
D_t2(28)=D_dia(8);
D_t2(33)=D_dia(9);
D_t2(37)=D_dia(10);
D_t2(40)=D_dia(11);
D_t2(42)=D_dia(12);
        [b,bint,r,rint,stats] = regress(D_t2,A_total);
        loss_r1=stats(4);
end

function x=best_group_m(C1,D1,J,C2,D2,m,D_t1,group_b,top_n,nn)
warning('off')
 x={};
rand_D={};
n=length(J);
if m==1
    s_size=2000;
end
if m==2
    s_size=2000;
end
if m==3
    s_size=2000;
end
        for uu=1:s_size
            aa=0.2+rand(n*(n+1),1);
            D_t=D_t1.*aa;
            D_t=D_t/sum(sum(D_t))*26;
            rand_D{uu}=D_t;      
        end
L={};
ss1=[];
ss2=[];
ss3=[];


   for i=1:nn
       if ~isempty(find(group_b==i))
           ss1(i)=1;
           ss2(i)=1;
           ss3(i)=1;
           continue;
       end
       group_now=[group_b,i];
       [A1,B1]=materix_trans(C1,D1,J);
       [A2,B2]=materix_trans(C2,D2,J);
       
%        A1=A1./sum(A1);
%        A2=A2./sum(A2);
%        
       A_total=[];
       A_diff=[A1(:,group_now),zeros(n*(n+1)/2,length(group_now));zeros(n*(n+1)/2,length(group_now)),A2(:,group_now)];
       A1(:,group_now)=[];
       A2(:,group_now)=[];
       A1_p=A1;
        A2_p=A2;
        A_total=[A_diff,[A1_p;A2_p]];

        ss_all=[];
        for uu=1:s_size
            D_t=rand_D{uu};
            [b,bint,r,rint,stats] = regress(D_t,A_total);
            ss_all(uu)=stats(4);          
        end
        
        ss_all=sort(ss_all);        
        ss1(i)=mean(ss_all(1:1));
%         ss2=mean(ss_all(1:20));
%         ss3=mean(ss_all(1:50));
        ss2(i)=mean(ss_all(1:20));
        ss3(i)=mean(ss_all(1:20));                   
        L{i}=group_now;
   end
   LL=[];
   for i=1:nn
       LL(i,1)=i;
       LL(i,2)=ss2(i);
   end
   A1=sortrows(LL,2);
   
   for i=1:top_n
       x{i}=sort(L{A1(i,1)});
   end
end

function x=optimum_x1(C1,C2,n,J,lb,ub,initialpopulation,s_size)
fun = @(x)loss_f(x,C1,C2,n,J);

nvars=n*(n+1);
A=[];
b=[];
Aeq=[];
beq=[];
nonlcon = @unitdisk;

%initialpopulation=lb+rand(n*(n+1),s_size).*(ub-lb);

options=optimoptions('ga','PopulationSize',s_size,'InitialPopulationMatrix',initialpopulation','MaxStallGenerations',500,'MaxGenerations',420000,'FunctionTolerance',1e-10);

x=ga(fun,nvars,A,b,Aeq,beq,lb,ub,nonlcon,options);


end

function x=optim_second(C1,C2,n,J,lb,ub,x0)
fun = @(x)loss_f(x,C1,C2,n,J);
nonlcon = @unitdisk;
A = [];
b = [];
Aeq = [];
beq = [];

%%options = optimoptions('fmincon','Display','iter','Algorithm','sqp');

x = fmincon(fun,x0,A,b,Aeq,beq,lb,ub,nonlcon) ;
end

function [J1,J2,diff_J]=J_cal_x(x,C1,C2,J,n)

x1=x(1:n*(n+1)/2);
x2=x((n*(n+1)/2+1):n*(n+1));
D_x1=zeros(n);
D_x2=zeros(n);
D_index=zeros(n);
l=1;
for k=1:n*(n+1)/2
    i=ceil(l/n);
    j=mod(l,n);
    if j==0
        j=n;
    end
    if i==0
        i=n;
    end
    while D_index(i,j)==1
         l=l+1;
         i=ceil(l/n);
         j=mod(l,n);
    if j==0
        j=n;
    end
    if i==0
        i=n;
    end
    end
    i=ceil(l/n);
    j=mod(l,n);
    if j==0
        j=n;
    end
    if i==0
        i=n;
    end

    D_x1(i,j)=x1(k);
    D_x2(i,j)=x2(k);
    D_x1(j,i)=x1(k);
    D_x2(j,i)=x2(k);
    D_index(i,j)=1;
    D_index(j,i)=1;
end

J1=calculateJ(C1,D_x1,J);
J2=calculateJ(C2,D_x2,J);

diff_J=zeros(n);
for i=1:n
    for j=1:n
        if J1(i,j)==0
            continue;
        end
        diff_J(i,j)=(J1(i,j)-J2(i,j))/(J1(i,j)+J2(i,j));
    end
end
end

function x=optimum_x(C1,C2,n,J,lb,ub,initialswarm_m,s_size)
fun = @(x)loss_f(x,C1,C2,n,J);

options = optimoptions('particleswarm','InitialSwarmMatrix',initialswarm_m','UseParallel',true,'SwarmSize',s_size,'MaxStallIterations',200,'MaxIterations',10000,'FunctionTolerance',0.0000001);

rng default  % For reproducibility
nvars = n*(n+1);
x = particleswarm(fun,nvars,lb,ub,options);

end

function loss_f_=loss_f(x,C1,C2,n,J)

x1=x(1:n*(n+1)/2);
x2=x((n*(n+1)/2+1):n*(n+1));
D_x1=zeros(n);
D_x2=zeros(n);

D_index=zeros(n);

l=1;
for k=1:n*(n+1)/2
    i=ceil(l/n);
    j=mod(l,n);
        if j==0
        j=n;
    end
    if i==0
        i=n;
    end
    while D_index(i,j)==1
         l=l+1;
         i=ceil(l/n);
         j=mod(l,n);
             if j==0
        j=n;
    end
    if i==0
        i=n;
    end
    end
    i=ceil(l/n);
    j=mod(l,n);
    if j==0
        j=n;
    end
    if i==0
        i=n;
    end

    D_x1(i,j)=x1(k);
    D_x2(i,j)=x2(k);
    D_x1(j,i)=x1(k);
    D_x2(j,i)=x2(k);
    D_index(i,j)=1;
    D_index(j,i)=1;
end

J1=calculateJ(C1,D_x1,J);
J2=calculateJ(C2,D_x2,J);


ee=0.00000001;
p=0.2;
loss_f_=0;
for i=1:n
    for j=1:n
        if (J1(i,j)==0 && J2(i,j)==0)
            continue;
        end
        l=(((J1(i,j)-J2(i,j))/(J1(i,j)+J2(i,j)))^2+ee)^(p/2);
        loss_f_=loss_f_+l;
    end
end

end

function Cal_J=calculateJ(C,D,J)

n=length(J);
J_structure=ones(n);
for i=1:n
    for j=1:n
        if J(i,j)==0
            J_structure(i,j)=0;
        end
    end
end

A=zeros(n*(n+1)/2,n*n);
X_zeros=find(J_structure'==0);


k=1;
for i=1:n
    for j=i:n
        B(k)=D(i,j);
        for l=1:n
            A(k,n*(i-1)+l)=A(k,n*(i-1)+l)+C(l,j);
            A(k,n*(j-1)+l)=A(k,n*(j-1)+l)+C(i,l);
        end
        k=k+1;
    end
end

J_aim=reshape(J',[n*n,1]);
A(:,X_zeros)=[];
J_aim(X_zeros)=[];
J_vec=A\B';

k=1;
J_cal=zeros(n);
for i=1:n
    for j=1:n
        if isempty(find(X_zeros==((i-1)*n+j)))
            J_cal(i,j)=J_vec(k);
            k=k+1;
        end
    end
end


m=1;

Cal_J=J_cal;


end

function [A,B]=materix_trans(C,D,J)

n=length(J);
J_structure=ones(n);
for i=1:n
    for j=1:n
        if J(i,j)==0
            J_structure(i,j)=0;
        end
    end
end

A=zeros(n*(n+1)/2,n*n);
X_zeros=find(J_structure'==0);


k=1;
for i=1:n
    for j=i:n
        B(k)=D(i,j);
        for l=1:n
            A(k,n*(i-1)+l)=A(k,n*(i-1)+l)+C(l,j);
            A(k,n*(j-1)+l)=A(k,n*(j-1)+l)+C(i,l);
        end
        k=k+1;
    end
end


A(:,X_zeros)=[];


end

function [c,ceq] = unitdisk(x)
c = x(1)^2 + x(2)^2 - 5;
ceq = [];
end
```
